# CRISPR-based environmental biosurveillance assisted via artificial intelligence design of guide-RNAs

**DOI:** 10.1101/2024.12.10.627849

**Authors:** Benjamín DuránVinet, Jo-Ann L. Stanton, Gert-Jan Jeunen, Ulla von Ammon, Jackson Treece, Xavier Pochon, Anastasija Zaiko, Neil J. Gemmell

**Affiliations:** Department of Anatomy, School of Biomedical Sciences, University of Otago, Dunedin, 9016, New Zealand; Department of Marine Sciences, University of Otago, Dunedin, 9054, New Zealand; Cawthron Institute, Nelson 7010, New Zealand; Institute of Marine Science, University of Auckland, Auckland 1142, New Zealand; Sequench, Nelson 7010, New Zealand

**Keywords:** Environmental DNA, Biosecurity, invasive species, CRISPR-Cas13a, neural network

## Abstract

Environmental biosecurity challenges are worsening for aquatic ecosystems as climate change and increased anthropogenic pressures facilitate the spread of invasive species, thereby broadly impacting ecosystem composition, functioning, and services. Environmental DNA (eDNA) has transformed traditional biomonitoring through detection of trace DNA fragments left by organisms in their surroundings, primarily by application of the quantitative polymerase chain reaction (qPCR). However, qPCR presents challenges, including limited portability, reliance on precise thermal cycling, and susceptibility to inhibitors. To address these challenges and enable field-deployable monitoring, isothermal amplification techniques such as Recombinase Polymerase Amplification (RPA) paired with Clustered Regularly Interspaced Short Palindromic Repeats and associated proteins (CRISPR-Cas) have been proposed as alternatives. We report here the development of CORSAIR (**<u>C</u>**RISPR-based envir**<u>O</u>**nmental biosu**<u>R</u>**veillance a**<u>S</u>**sisted via **<u>A</u>**rtificial **I**ntelligence guide-**<u>R</u>**NAs), that harnesses the programmability of the CRISPR-Cas technology, RPA and the artificial intelligence (AI)-based tool Activity-informed Design with All-inclusive Patrolling of Targets (ADAPT) to deploy a swift RPA-CRISPR-Cas13a-based method that detects eDNA from two invasive species as proof of concept: *Sabella spallanzanii* and *Undaria pinnatifida*. CORSAIR showcased a robust, streamlined method augmented by ADAPT, reaching a high specificity when tested against co-occurring species and a 100% agreement with 12 PCR-benchmarked eDNA samples, reaching a sensitivity of 0.34 copies uL^-1^ in 1 hour with a cost of 3.5 USD per sample; thus highlighting CORSAIR as a powerful environmental biosurveillance platform for environmental nucleic acid detection.

**Graphical abstract.**
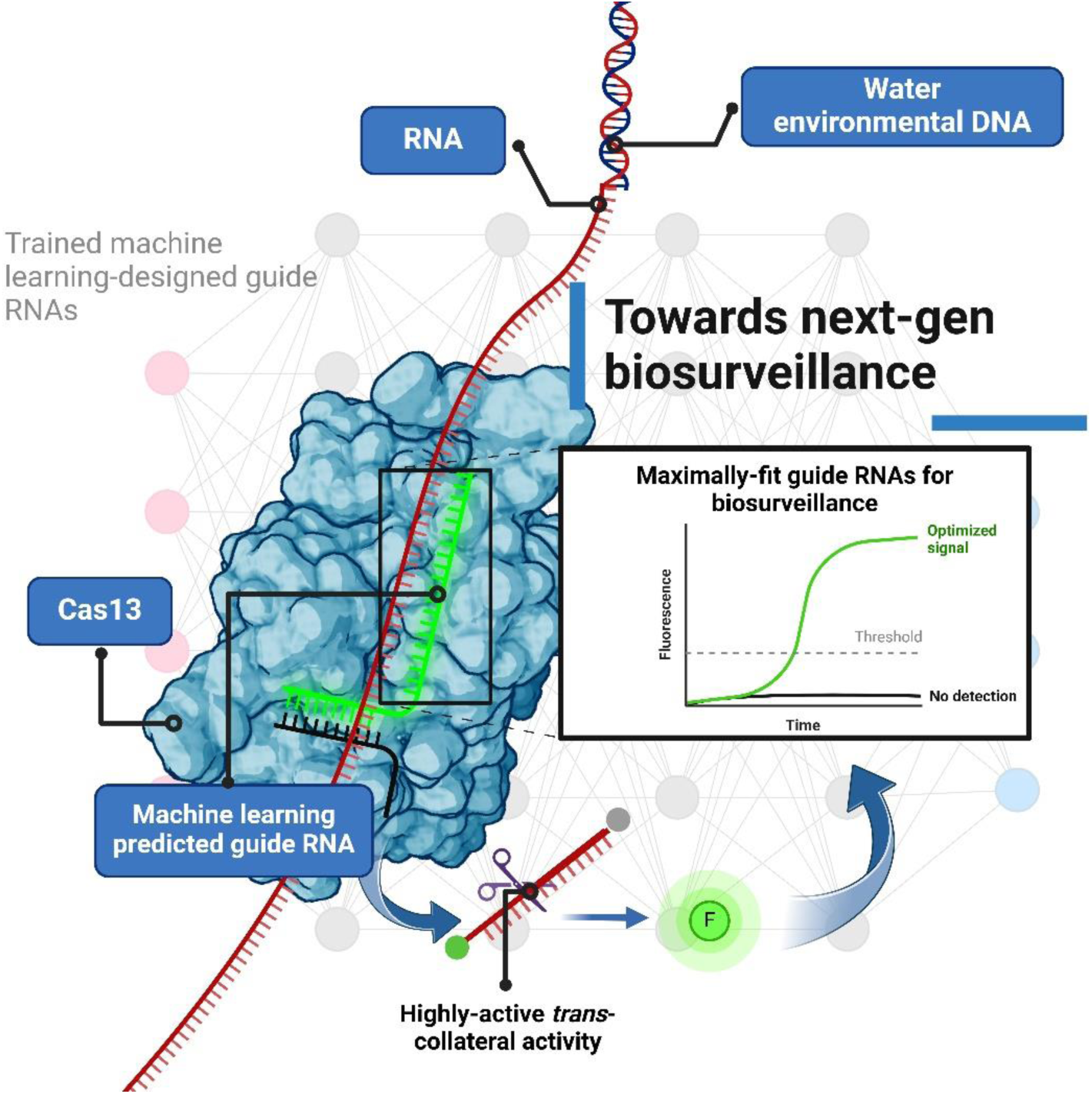

## Introduction

Environmental biosecurity challenges in aquatic ecosystems are evolving under pressures from climate change, invasive species, and human activities, which collectively affect ecosystem sustainability, endemic biodiversity, and related economic activities (1–3). Combatting these pressing issues demands innovative, reliable, and robust solutions to enhance biosurveillance efforts to manage and mitigate biosecurity risks (4, 5). Current biosurveillance efforts are mostly based on traditional biomonitoring (e.g. capture), which can be laborious and require morphological taxonomy expertise (6, 7). Recent studies have highlighted environmental DNA (eDNA) as a novel, minimally invasive approach for the precise and sensitive detection of bioinvasive species. By targeting specific gene regions, molecular tools such as polymerase chain reaction (PCR) allow ecologists to efficiently detect the presence or absence of species of interest at scale (8–10). Although PCR-based methods now have portable options (11, 12), they still require precise temperature cycling and may be susceptible to PCR inhibitors (13).

A proposed solution is using isothermal amplification technologies for eDNA detection to tackle current PCR challenges, however, the most commonly used isothermal amplification technologies have critical shortcomings that make initial testing a complex task. For example, recombinase polymerase amplification (RPA) provides quick DNA amplification, it lacks specificity leading to false negatives and high background noise, while loop-mediated isothermal amplification (LAMP) approaches display high specificity but has a complex primer design, which can lead to false-negatives and undesired poor sensitivity performance (14, 15).

Recent studies are harnessing the natural prokaryotic adaptive immune system known as CRISPR- Cas technology (Clustered regularly interspaced short palindromic repeats and associated CRISPR proteins) (16, 17), coupled with isothermal methods such as RPA and LAMP (14, 18), to enrich the sample and circumvent standalone drawbacks, further increasing the specificity and sensitivity of the overall diagnostic platform, ultimately detecting eDNA isothermally with promising results, including high specificity, sensitivity, and robustness while also enabling the possibility of point-of-need in field testing if required (19–21). Moreover, CRISPR technology can be applied directly to briefly heated, unprocessed saliva or urine samples, demonstrating its effectiveness even in the presence of residual PCR inhibitors, while remaining suitable for field deployment through lateral flow screening (22).

In short, CRISPR-Cas technology harnesses RNA molecules known as CRISPR RNAs (crRNAs) that guide the Cas nucleases to the target by nucleotide complementarity. Specifically, Cas13a, upon recognizing and cleaving its RNA target (*cis* activity), triggers a *trans*-collateral cleavage activity that further targets short RNA molecules. This property has been harnessed to develop rapid diagnostic platforms by coupling short RNAs with a fluorophore (e.g., 6-carboxyfluorescein – 6-FAM), biotin for lateral-flow screening, and a quencher (e.g., Black hole quencher 1 – BHQ1). A positive reaction produces either a fluorescence peak or a band (23–25)

Our framework proposes an integrated approach combining CRISPR, RPA and artificial intelligence (AI) to enhance environmental biosurveillance efforts using eDNA, namely **<u>C</u>**RISPR-based envir**<u>O</u>**nmental biosu**<u>R</u>**veillance a**<u>S</u>**sisted via **<u>A</u>**rtificial **I**ntelligence guide-**<u>R</u>**NAs (CORSAIR, Figure 1). CORSAIR harnesses different cutting-edge technologies, starting with trained AI modelling for the discovery of highly active guide-target pairs, specifically, RPA forward and reverse primers, and a crRNA target (Step 1, Figure 1), to then use the RPA-T7-CRISPR-LwaCas13a designs for isothermal detection of targets of interest (Steps 2-4, Figure 1).

**Figure 1.**
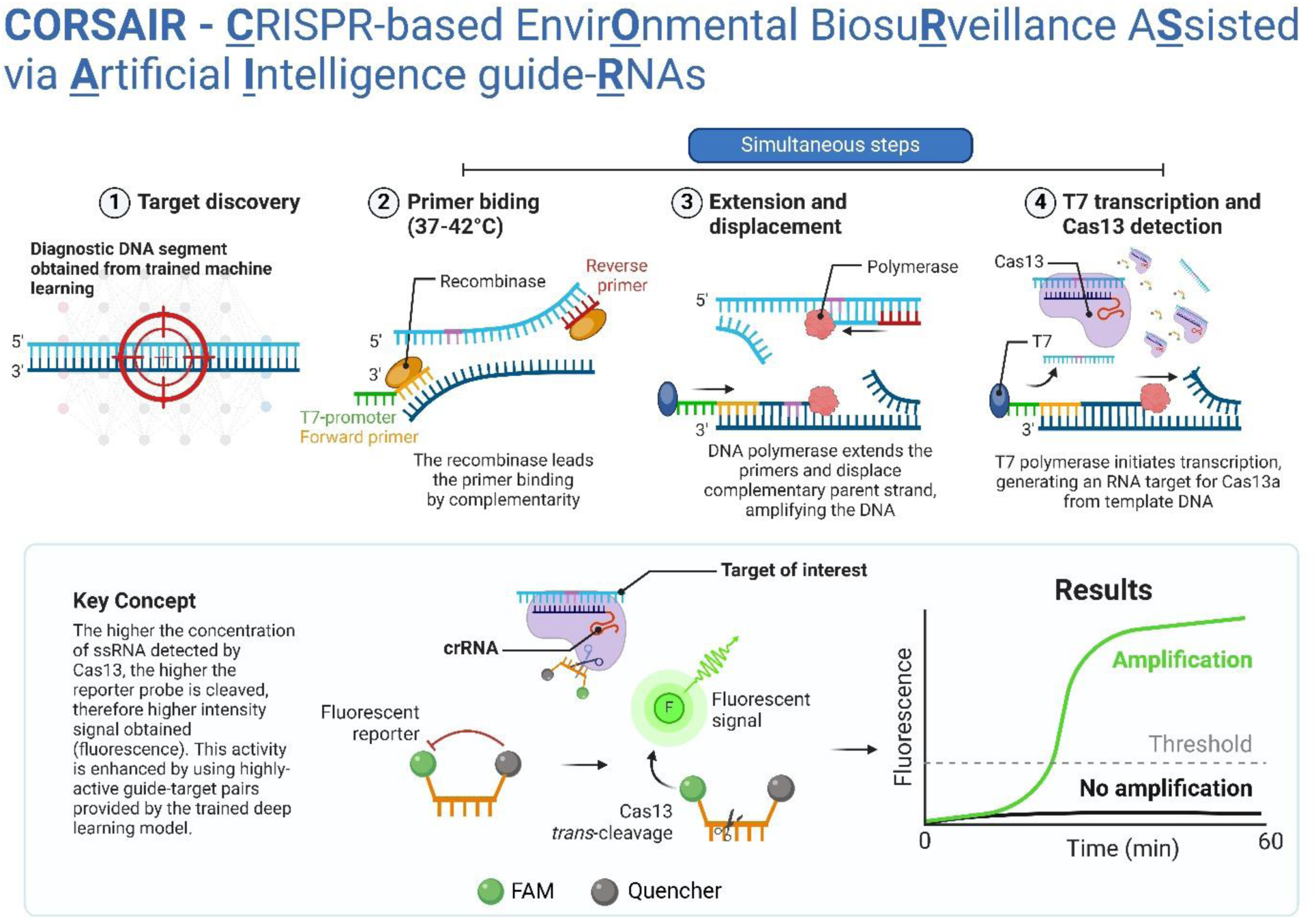
CORSAIR platform concept. (1) The best diagnostic segment is discovered using ADAPT. (2) Using the guide-target pairs provided, the target is amplified (3), transcribed and detected by LwaCas13a (4). The key concept of CRISPR-eBx technology is that as more target concentration is cleaved, reporter probes will also be cleaved, providing a fluorescence signal. This activity can be currently optimized by using trained artificial intelligence models. ADAPT: Activity-informed Design with All-inclusive Patrolling of Targets; CRISPR-eBx: CRISPR-based environmental biosurveillance.

In this study, we focused on designing crRNAs leveraging a trained, end-to-end convolutional neural network termed ADAPT (Activity-informed Design with All-inclusive Patrolling of Targets; https://adapt.run/) that facilitates the identification of optimal, highly active diagnostic crRNA configurations providing a ranked list of guide-target pairs (26). This is one of the main bottlenecks of CRISPR technology since different crRNA configurations on the spacer region (complementarity region) directly impact the on-target and off-target activity of Cas nucleases (27). Thereby, ADAPT maximizes on-target specificity and sensitivity and minimizes unwanted off-target activity, delivering highly active and precise diagnostic assays for *Leptotrichia wadeii* Cas13a nuclease (LwaCas13a) (28), that we used for CRISPR-based environmental biosurveillance (CRISPR-eBx).

We used ADAPT on two invasive species in New Zealand as proof-of-concept, *Sabella spallanzanii* (Mediterranean fanworm, (29)) and *Undaria pinnatifida* (Asian seaweed, (30)), for which we designed species-specific CRISPR-eBx assays targeting their *cytochrome c oxidase subunit I* (COI) gene. We then evaluated CORSAIR under various controlled conditions to optimize the detection platform, followed by testing on *S. spallanzanii* and *U. pinnatifida* eDNA samples previously validated with qPCR. CORSAIR demonstrated 100% concordance with PCR-based methods and represents a streamlined CRISPR-eBx platform, enabling rapid and efficient target detection for biomonitoring known and emerging environmental biosecurity threats using LwaCas13a.

## Material and Methods

### Construction of the target reference

All *in silico* analyses were performed in Geneious Prime (2024.0.5; https://www.geneious.com). We used COI as the reference gene to build a consensus target for our CORSAIR assay. We retrieved *Sabella spallanzanii* (KY472787.1) and *Undaria pinnatifida* (NC_023354.1) COI genes from GenBank (release 258.0; www.ncbi.nlm.nih.gov/genbank/). Each reference sequence was used in GenBank megaBLAST (version 2.14.0; (31)) to retrieve all sequences with a ≥99.6% and ≥99.5% pairwise identity (Supplemental Table 1), obtaining 18 and 14 sequences, respectively. Accession numbers are available in Supplemental Table 1.

Retrieved sequences for each species were aligned using the Clustal Omega alignment algorithm (version 1.2.2; Supplemental File 1; (32)). High-coverage regions and polymorphic sites were identified with a 50% minimum sequence frequency threshold. Then, we generated consensus sequences for *S. spallanzanii* and *U. pinnatifida* with a 0% nucleotide call threshold (Supplemental Table 2) and used them in the next step of the ADAPT pipeline. We also used these consensus sequences for gene synthesis and custom plasmid construction (GenScript, USA, custom product).

### ADAPT local deployment for environmental DNA target discovery

ADAPT (version 1.6.0; https://adapt.run/) is an end-to-end deep-learning neural network that predicts highly active CRISPR RNAs (crRNAs) for viral diagnostic deployment, trained with 19,209 guide-target pairs outcomes from *Leptotrichia wadeii* Cas13a (LwaCas13a) to enhance *trans* collateral cleavage activity (26). Guide-target pairs refer to the spacer region of the crRNA and flanking primers for RPA (26). We used ADAPT with the COI consensus sequences (*S. spallanzanii* and *U. pinnatifida*) as “on-target sequences” in FASTA format, while incorporating identified off-target sequences as “excluded sequences” in FASTA format. Additional parameters, including primer length, spacer length, spacer specificity (Supplemental Table 3), were applied to customize and optimize LwaCas13a crRNA modelling and diagnostic performance prediction. The final command used for *S. spallanzanii* and *U. pinnatifida* is provided in Supplemental Table 3. The output of ADAPT was a .TSV file with the predicted activity, crRNA spacer sequence (28 nucleotides) and RPA primer sequences (30 nucleotides each). Only crRNAs spacer sequences were checked with megaBLAST.

As previously reported, forward primers were appended with a T7 promoter sequence at the 5’ end (5’-gaaatTAATACGACTCACTATAGGG-3’) (33–35). All primers, crRNAs and probes were synthesized by GenScript US as custom products (Supplemental Table 4). The ADAPT repository, installation and running guide is publicly available on GitHub (https://github.com/broadinstitute/adapt). A summary of the overall workflow is illustrated in Figure 2.

**Figure 2.**
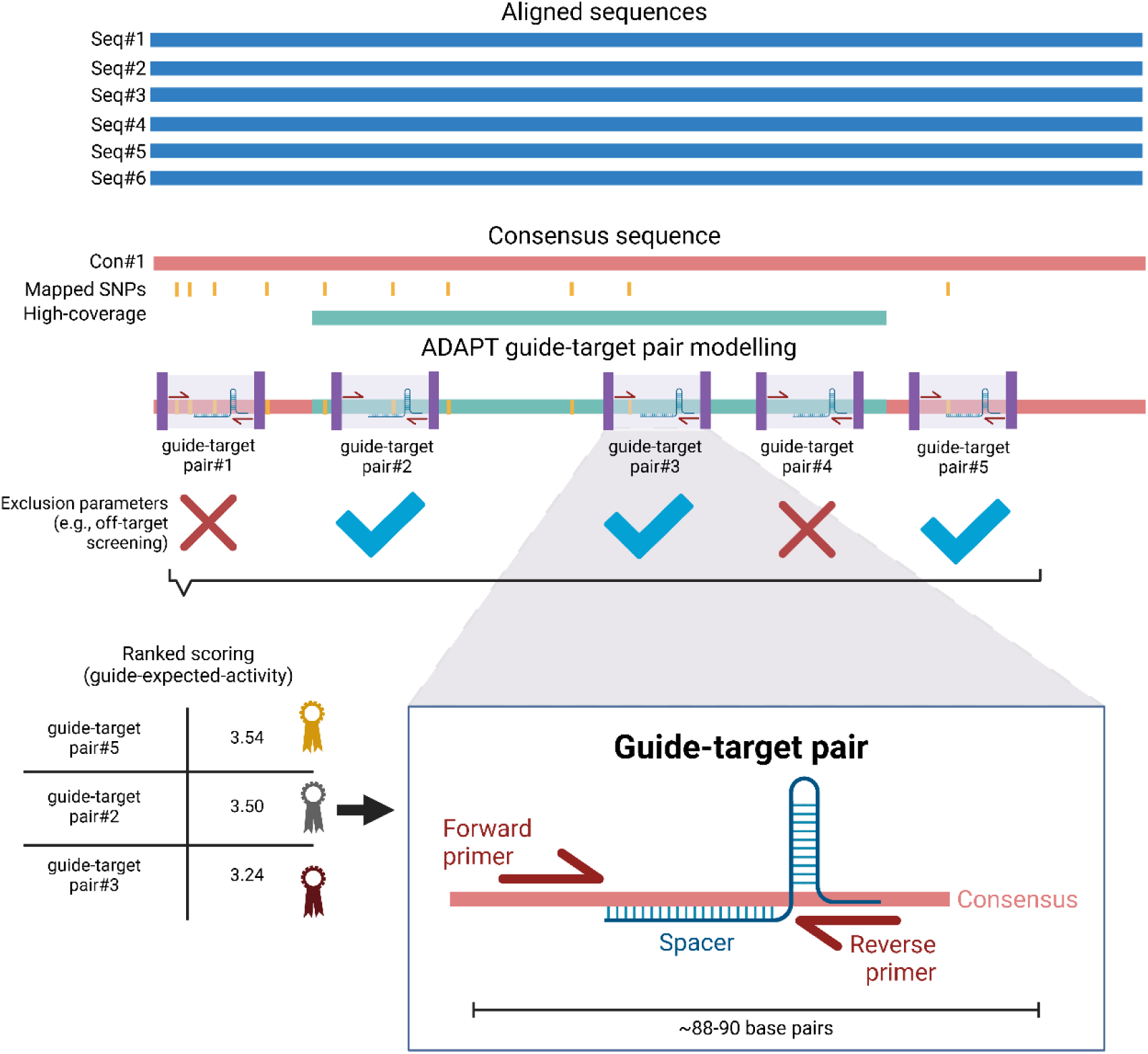
Simplified illustration of the overall process to obtain guide-target pairs. Sequences were aligned to map single nucleotide polymorphism (SNP) sites (yellow) and high-coverage regions (cyan). A consensus sequence is generated with all the previously obtained annotations (red). This consensus is used in ADAPT, where different guide-target pairs are obtained (purple squares). Another ADAPT run is performed with exclusion parameters (e.g., exclude off-targets and increase specificity) to filter guide-target pairs. Filtered guide target pairs are ranked and selected depending on their gRNA expected activity predicted by the ADAPT model. Each guide-target pair has a total length of ∼88-90 base pairs when considering forward/reverse primers and spacer regions.

### Genomic DNA extraction

Genomic DNA (gDNA) of *S. spallanzanii* was extracted using the DNeasy^®^ Qiagen Blood and Tissue (Qiagen, #69504) following the manufacturer instructions. Extracted gDNA samples were stored at - 20°C until further processing. We used a previously optimized qPCR assay to check gDNA signal, as previously reported (8). The qPCR configuration was as follows: 95 °C for 3 min, followed by 30 cycles at 95 °C for 15 s and 60 °C for 30 s. The primers were as following (forward primer: GCTCTTATTAGGCTCTGTGTTTG; reverse primer: CCTCTATGTCCAACTCCTCTTG; probe: AAATAGTTCATCCCGTCCCTGCCC).

*Undaria pinnatifida* gDNA extractions followed the PDQeX extraction method as previously reported (36). Fresh *U. pinnatifida* tissue samples were collected with a 3 mm hole punch (∼10 mg), the tissue rinsed with Milli-Q water and manually homogenized with 80 uL 1X Green Buffer using a pestle. 40 uL of the homogenate was added to the extraction mastermix (48 uL of RNAse-free water, 40 uL homogenate, 10 uL 10X Green Buffer, 2 uL PrepGem enzyme). The extraction mastermix was dispensed into PDQeX extractor cartridges and run through the following protocol in the PDQeX 2400 device: 37°C for 5 min, 75°C for 5 min and 95°C for 2 min. At the end of the program, extracts containing purified DNA were collected in 0.2-ml PCR tubes. Purified DNA samples were stored in 0.2-ml PCR tubes at −20°C until further processing. *U. pinnatifida* gDNA and eDNA samples were checked with qPCR (95°C for 3 min, then cycled 45 times: 95°C for 15 s, 61°C for 15 s and 72°C) as previously described (37), using the following primers (forward primer: TACAGCAATGTCTGTTTTTATCC; reverse primer: ACATTATACAACTGATGATTTCCC; probe: ATTGCAATTAGCTAGCCCTG).

Both gDNA extractions were quantified with a Qubit 4 Fluorometer (ThermoFisher Scientific) following manufacturer instructions. The initial concentration for *S. spallanzanii* gDNA was 18 ng µL^-1^ and *U. pinnatifida* gDNA was 3.4 ng µL^-1^. Consecutive ten-fold dilutions were set up and used immediately for gDNA sensitivity assays.

### Environmental DNA samples

Processing of environmental DNA samples were conducted in a dedicated PCR-free laboratory. Benches and equipment were decontaminated using a 10-min exposure to 10% bleach solution and wiped with ultrapure water (UltraPure™ DNase/RNase-Free Distilled Water, Invitrogen™) before laboratory work commenced. As CORSAIR also works with RNA, RNaseZAP™ (Invitrogen) was used to remove RNAses from benches and equipment.

Environmental samples from *S. spallanzanii* infected sites were obtained from von Ammon et al. (38). Environmental samples from *U. pinnatifida* containing sites (Otago Harbour – OH#1–3) were obtained following the protocol previously described in Jeunen et al. (39). OH2 and OH3 eDNA samples were obtained by filtrating 1L seawater while OH1 was obtained by filtrating 2L seawater. We used a 1.2 µm ∼30 mm cellulose acetate filter (Whatman) for OH1 and OH3 eDNA samples, while OH2 used a 0.22 µm ∼30 mm cellulose acetate filter (Whatman). It has been reported that either 1L or 2L works for eDNA capture, purification and detection (40). Lastly, environmental samples free of *S. spallanzanii* and *U. pinnatifida* were obtained from Doubtful Sound, Fiordland, New Zealand, and were collected as part of a previous study (41). All samples were aliquoted in the PCR-free clean room and always stored at −20°C. Aliquots were taken out of the PCR-free clean room and used for qPCR or CORSAIR assays.

### Cas13 fluorescence assays

The Cas13 mastermix for fluorescence assays (Supplemental Table 5) was standardized and modified from previously reported Cas13-based fluorescence assays (28, 33–35, 42–44). For all CORSAIR reactions, we used a 1:1 (LwaCas13a:crRNA) ratio. The mastermix composition is as follows: 5.18 µL of RNase-free water (Invitrogen, #10977015), 4 µL of 5X optimized Cas13a reaction buffer (see Supplemental Table 6) or 2 µL 10X Cas13a reaction buffer (GenScript, #Z03486), 0.5 µL of Murine RNAse inhibitor (New England Biolabs - NEB, #M0314S), 0.8 µL of ribonucleotide mix (NEB, #N0466S), 1 µL of 10 µM reporter (poly-U_5_ or poly-U_15_, GenScript, custom), 0.96 µL of 10 µM forward primer (GenScript, custom), 0.96 µL of 10 µM reverse primer (GenScript, custom), 0.6 µL of T7 RNA polymerase (NEB, #M0251S), 2 µL of 500 nM LwaCas13a (GenScript, #Z03486; diluted in RNAse-free water and 1X RNAse-free PBS pH 7.4, Invitrogen, #AM9624), 1 µL of 1 µM crRNA (GenScript, custom) and 1 µL of 280 mM MgOAc (TwistDx). All components were added in the described order. We considered a 5% pipetting error when scaling up the reactions. All mastermixes were freshly produced and used immediately. Cas13a optimized reaction buffer (Supplemental Table 6) composition was used as previously reported (45).

After setting up the mastermix, it was slowly mixed with one RPA pellet (TwistDx). 18 uL of mastermix was aliquoted in a white 8-strip tube using a 96-well adaptor (Roche Diagnostics, LightCycler^®^ 8- Tube Strip Adapter Plate, #06612598001), and then 2 µL of sample added. Then, strips were spun down for 15 seconds at full speed in a minifuge and immediately run in LC480 II LightCycler qPCR systems (Roche Diagnostics) at 39°C as previously reported (46), for one hour with data acquisitions every 60 seconds in the FAM channel (excitation at λ465 nm and detection at λ510 nm). RNAse-free water was used as a negative control in all assays. A list of the samples used in this study is provided in the Supplemental Table 8. A cost breakdown is provided in Supplemental Table 9.

### Overall calculations, data plots and figures

One-way ANOVA test, t-test and correlation were performed with Prism 10 (Version 10.4.0 for Windows, GraphPad Software, Boston, Massachusetts USA, www.graphpad.com) using a significant value equal to p < 0.01 where applicable, unless otherwise indicated. All data plots (heatmap, column plots, and curves) were created with Prism 10. All illustrations were created with BioRender (www.biorender.com/). Fold-change calculations were done with end-point raw intensity data, dividing the control-subtracted end-point raw intensity from the target with the average of the end-point raw intensity of controls. To standardize the fold-change calculations, we used a fixed no-target raw intensity control average value (∼0.3) from 13 no-target control replicates across different assays.

## Results

### CRISPR RNA design with ADAPT for local biosurveillance deployment

ADAPT provided six and four highly active guide-target pairs (i.e., forward primer, crRNA and reverse primers, **Figure 2**) for *S. spallanzanii* and *U. pinnatifida,* respectively, with no off-target hits (using 100% hit search) on the spacer region when checked using megaBLAST. We selected the best and worst guide-target pair for each species to test further (Supplemental Table 4). A full list of guide-target pairs obtained from ADAPT with their scoring for *S. spallanzanii* and *U. pinnatifida* is provided in Supplemental Table 7.

### Optimized Cas13a buffer displays better diagnostic activity than the provided manufacturer buffer

To screen whether ADAPT designs worked as intended for the CORSAIR platform, we first tested a single guide-target pair for *U. pinnatifida* detection (U-cr180) against 10 pM of plasmid DNA using poly-U_5_ in two different conditions: a previously optimized buffer for Cas13a diagnostics (45) and the provided Cas13a reaction buffer from the manufacturer. We observed that the U-cr180 guide-target pair had a faster activation speed with the optimized Cas13a buffer rather than the manufacturer buffer (p = 0.03 when using p < 0.05, Figure 3). The optimized reaction buffer was 6 minutes faster than the proprietary buffer in getting beyond the arbitrary threshold (set later at 7- fold change units or ∼2.1 raw intensity). Additionally, the Cas13a optimized reaction buffer reached saturation point at ∼16 min (i.e., faster activation speed) while the proprietary buffer reached saturation at ∼39 min, making the Cas13a optimized reaction buffer ∼2.4 times faster, aligned with similar results from the study which described this optimized buffer (45). We understand activation speed as the elapsed time of LwaCas13a reaction to reach saturation when providing a positive result. Accordingly, we used the optimized Cas13a reaction buffer for all following CORSAIR assays.

**Figure 3.**
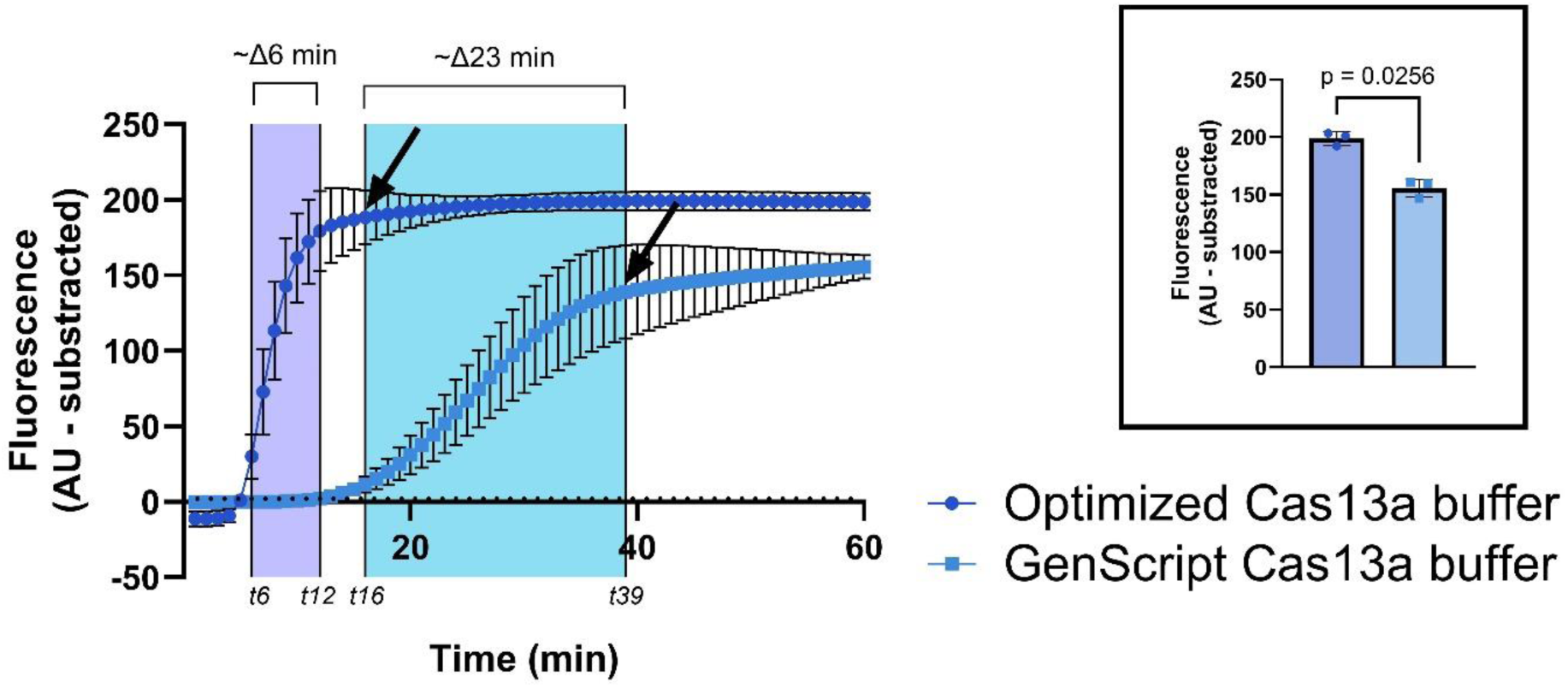
Cas13a diagnostic activity under two different buffers. The optimized Cas13a buffer provided a faster activation by minute 6 (t6) than the GenScript Cas13a buffer at minute 12 (t12); this difference is highlighted in a purple rectangle. Optimized Cas13a buffer also performed better, reaching saturation point faster by minute 16 (t16, black arrow) versus minute 39 (t39, black arrow) provided by the GenScript Cas13a buffer); this difference is highlighted in a cyan rectangle. The black outlined box showcases endpoint raw intensity significance between both groups, with optimized Cas13a having significantly more intensity (p = 0.02). Columns represent the mean, and error bars show ± SD (n = 3). The p-value was significant at p < 0.05.

### A shorter poly-U5 reporter displays better diagnostic activity than a longer poly-U15 reporter

We delved into testing a novel reporter for LwaCas13a (U_15_-reporter, 5’-6-FAM/UUU UUU UUU UUU UUU/BHQ1-3’; (47)) versus the traditional reporter (U_5_-reporter, 5’-6-FAM/UUU UU/BHQ1-3’; (23, 35)) to explore whether a shorter or longer poly-uracil influence the diagnostic performance of CORSAIR. We tested CORSAIR under the same reporter concentration (500 nM final), using a fixed plasmid concentration (10 pM or ∼0.3 ng/µL) containing the corresponding target sequence to set up the final mastermix composition for further CORSAIR assays **(Figure 3)**. The U_5_-reporter displayed significantly better end-point intensity fold change after 1h of reaction (p<0.001) across all crRNAs **(Figure 4A)**. Moreover, we observed that all crRNAs displayed similar activation speeds in the 10 pM range when using the Poly-U_5_ or Poly-U15 reporter **(Figure 4)**. Thus, poly-U5 was selected for all further assays.

**Figure 4.**
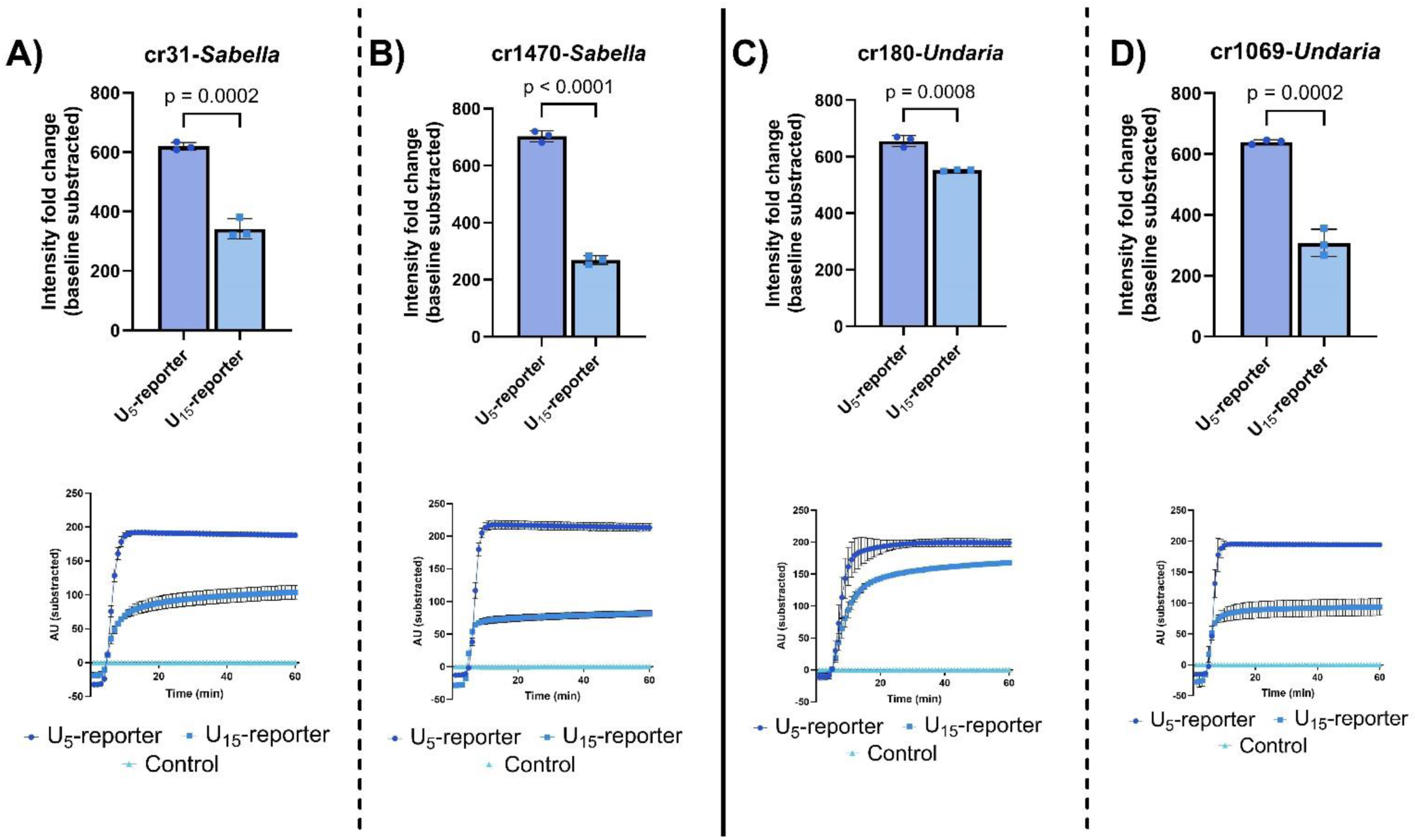
Poly-U_5_-reporter versus Poly-U_15_-reporter. The intensity fold change endpoint is shown in the upper panel, and the corresponding fluorescence plot of the reaction is shown in the bottom panel for **(A and B)** *Sabella spallanzanii* and **(C and D)** *Undaria pinnatifida*. All data points were obtained with n = 3. Columns represent the mean, and error bars show ± SD (n = 3). All p-values were significant (using p<0.01). Poly-U_5_: 5’-6-FAM/UUU UU/BHQ1-3’; Poly-U_15_: 5’-6-FAM/UUU UUU UUU UUU UUU/BHQ1-3’.

### ADAPT predicted scoring showed no difference between the best and worst guide-target pair when using gDNA

Additionally, the predicted score versus the end-point fold change output was only significant (p<0.01) for *S. spallanzanii* guide-target pairs S-cr31 vs S-cr1470, with p = 0.0035 for plasmid DNA tests when we used the Poly-U_5_ reporter, but these guide-target pairs were not significant when tested against gDNA (p = 0.5443). For *U. pinnatifida* guide-target pairs, U-cr180 versus U-cr1069 assays on plasmid DNA and gDNA were both not significant, with p = 0.2699 and p = 0.3281, respectively (**Supplemental Figure 1)**.

### Off-target assays with co-occurring marine species showcase CORSAIR as a specific assay for environmental DNA biosurveillance

To assess whether the CORSAIR assays were working as intended, we first tested internal off-target activity on plasmids containing the region of interest. Tests were performed on gDNA from *S. spallanzanii* and *U. pinnatifida* and human DNA (extracted from induced pluripotent stem cells provided by Dr Indranil Basik from the Department of Biochemistry, University of Otago) to assess detection potential and limitations. No visible off-target activity was observed across species after 1h **(Figure 5A),** in concordance with previous *in silico* results that showed no hits in NCBI public database. Guide-target pairs S-cr31 and U-cr180 for *S. spallanzanii* and *U. pinnatifida*, respectively, were selected for further experiments to reduce the number of assays to screen.

**Figure 5.**
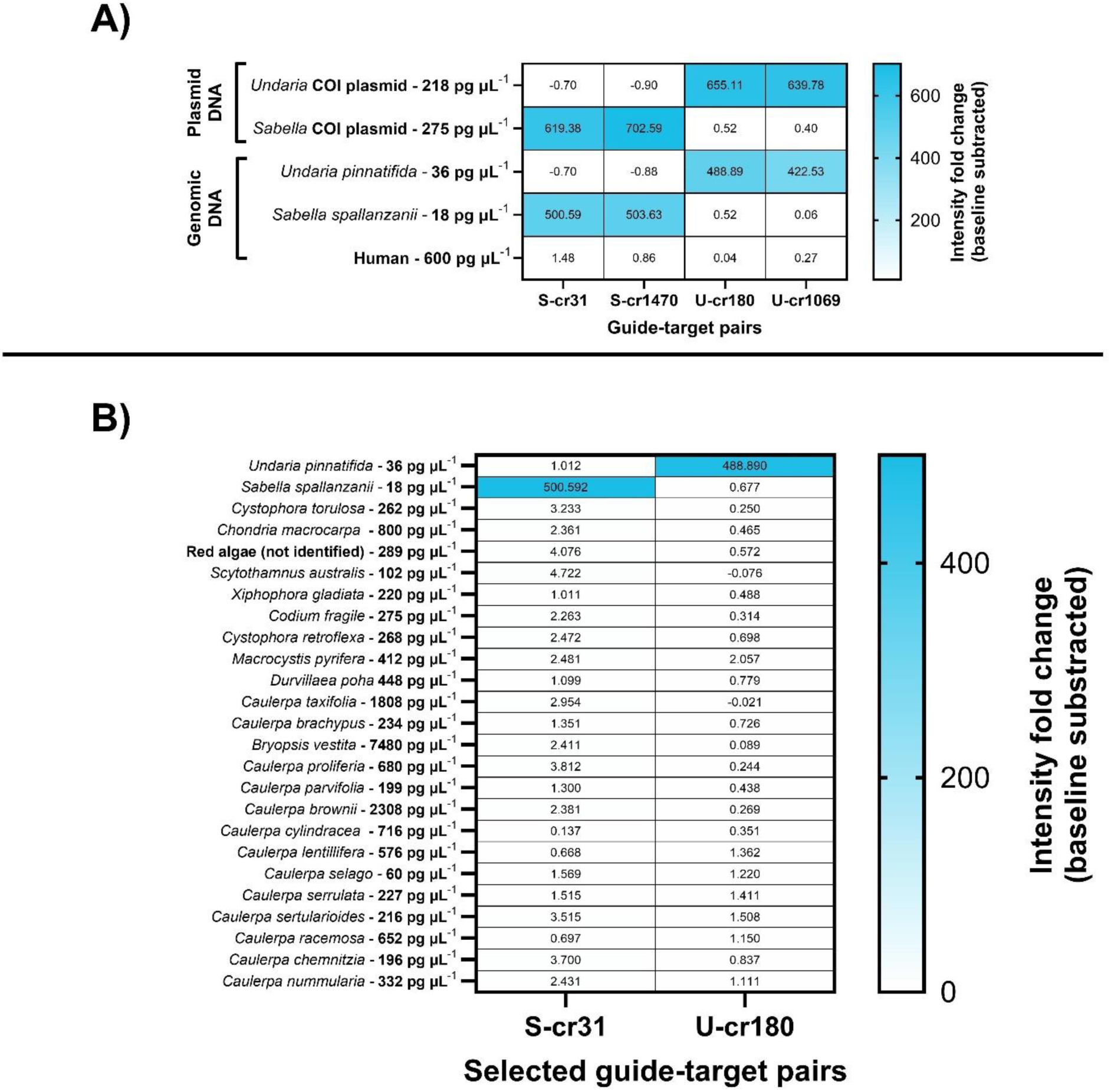
CORSAIR off-target assays with co-occurring species. All boxes represent the mean (n=3) of the end-point fold change after subtracting the baseline. **(A)** Internal off-target assays versus plasmid DNA containing *S. spallanzanii*/*U. pinnatifida* target inserts, gDNA and human gDNA. **(B)** Off-target activity against 23 other co-occurring marine species. CORSAIR was run for 1 hour. S: *Sabella spallanzanii*; U: *Undaria pinnatifida*; cr: CRISPR RNA; COI: Cytochrome c oxidase subunit I; CORSAIR: CRISPR-based environmental biosurveillance assisted via Artificial Intelligence guide-RNAs.

Next, we tested CORSAIR potential off-target activity in a more realistic off-target assay setup, using gDNA from 23 other marine species that could co-occur with our target species (Supplemental Table 8), ranging in tested concentration from 18 pg µL^-1^ to 7480 pg µL^-1^, which is a similar range from ∼4.3 nM to ∼10 pM **(Figure 5B)**. Additionally, to comprehensively reduce potential off-target effects and call positives, we set an arbitrary positivity threshold of 7-fold-change units as maximum background noise, similar to other CRISPR studies (44, 48, 49). The fold-change arbitrary threshold was obtained by multiplying five times the standard deviation of the average end-point raw intensity from no-target controls divided by their average end-point raw intensity, providing ∼2.1 points of raw fluorescence intensity equal to ∼7-fold-change units.

### CORSAIR is a highly sensitive biosurveillance tool with a semi-quantitative window

We challenged CORSAIR assay sensitivity in plasmid DNA and gDNA, obtaining attomolar sensitivity with a significant endpoint fold-change (p < 0.05) when compared with control (**Supplemental Figure 2**). The tested guide-target pairs obtained a theoretically semi-quantitative approach in plasmid DNA (R^2^=0.84 for *S. spallanzanii* and R^2^=0.85 for *U. pinnatifida,* **Figure 6A**) and in gDNA for both crRNAs tested (S-cr31 and U-cr180) when testing in high, medium-high, low-high, medium and low concentrations, obtaining R^2^ values equal to 0.77 and 0.81, respectively (**Figure 6B)**. S-cr31 and U-cr180 showcased a sensitivity down to 10 aM in plasmid DNA (equal to single copy ranges, ∼ 5 copies µL^-1^). Moreover, S-cr31 and U-cr180 achieved gDNA detection down to 0.018 pg µL^-1^ and 0.036 pg µL^-1^, respectively.

**Figure 6.**
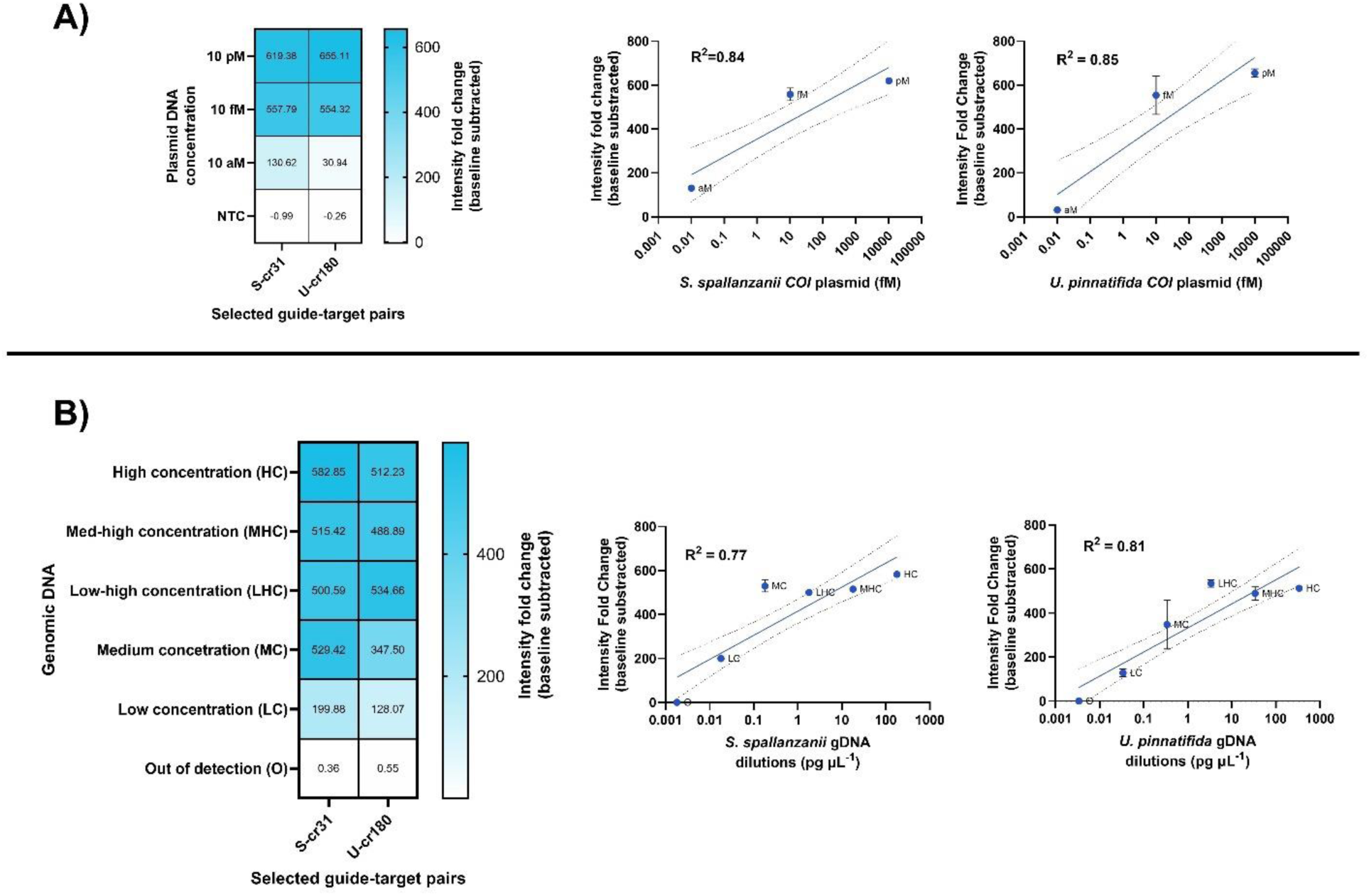
CORSAIR sensitivity and semi-quantitative windows. All boxes represent the mean (n = 3) **(A)** Sensitivity in over plasmid constructs. **(B)** Sensitivity over genomic DNA of target species. Data points represent the mean, and error bars show ± SD (n = 3). The dotted line highlights a 95% confidence interval. S: *Sabella spallanzanii*; U: *Undaria pinnatifida*; cr: CRISPR RNA; COI: Cytochrome c oxidase subunit I; CORSAIR: CRISPR-based environmental biosurveillance assisted via Artificial Intelligence guide-RNAs.

### Environmental DNA assays showcase CORSAIR as a specific and highly sensitive biosurveillance tool

To finalize the trial of our newly developed CORSAIR assays, we tested different eDNA samples that were known to contain *S. spallanzanii* (Marsden Cove), *U. pinnatifida* (Otago Harbour), as well as a control site where neither species were present (Doubtful Sound). All eDNA samples had been previously benchmarked using species-specific qPCR or ddPCR from another study (38) to give a point of comparison for CORSAIR. The results obtained from CORSAIR were highly sensitive and species-specific **(Figure 7).** Briefly, comparing the qPCR results from Otago Harbour eDNA samples (n = 3), we found that all were positive for qPCR and CORSAIR for *U. pinnatifida* (100% positive agreement), achieving a detection down to 59.2 copies µL^-1^. A similar result was obtained from the Marsden Cove eDNA samples (n = 6), which were benchmarked with ddPCR for *S. spallanzanii* and *U. pinnatifida*. All tested samples were ddPCR and CORSAIR positive for *S. spallanzanii* (100% positive agreement) and all negative for *U. pinnatifida* (100% negative agreement). Detection sensitivity was down to 0.34 copies µL^-1^ when detecting *S. spallanzanii* eDNA. All positives exceeded the arbitrary positivity calling threshold set earlier in previous assays in this study (. Moreover, when ddPCR and qPCR copies uL^-1^ were fit into a curve, *Sabella* eDNA samples showcased an R^2^ = 0.84, while *Undaria* eDNA samples had an R^2^ = 0.53, respectively (Supplemental Figure 3). We further explored CORSAIR assays off-target, testing eDNA samples from Doubtful Sound taken at different depths (0m – D0, 5m – D5, and 500m – D500), where no known invasion of *S. spallanzanii* or *U. pinnatifida* have been reported. All samples were negative for qPCR and CORSAIR. The raw fluorescence intensity plot of all positive eDNA samples are provided at Supplemental Figure 4.

**Figure 7.**
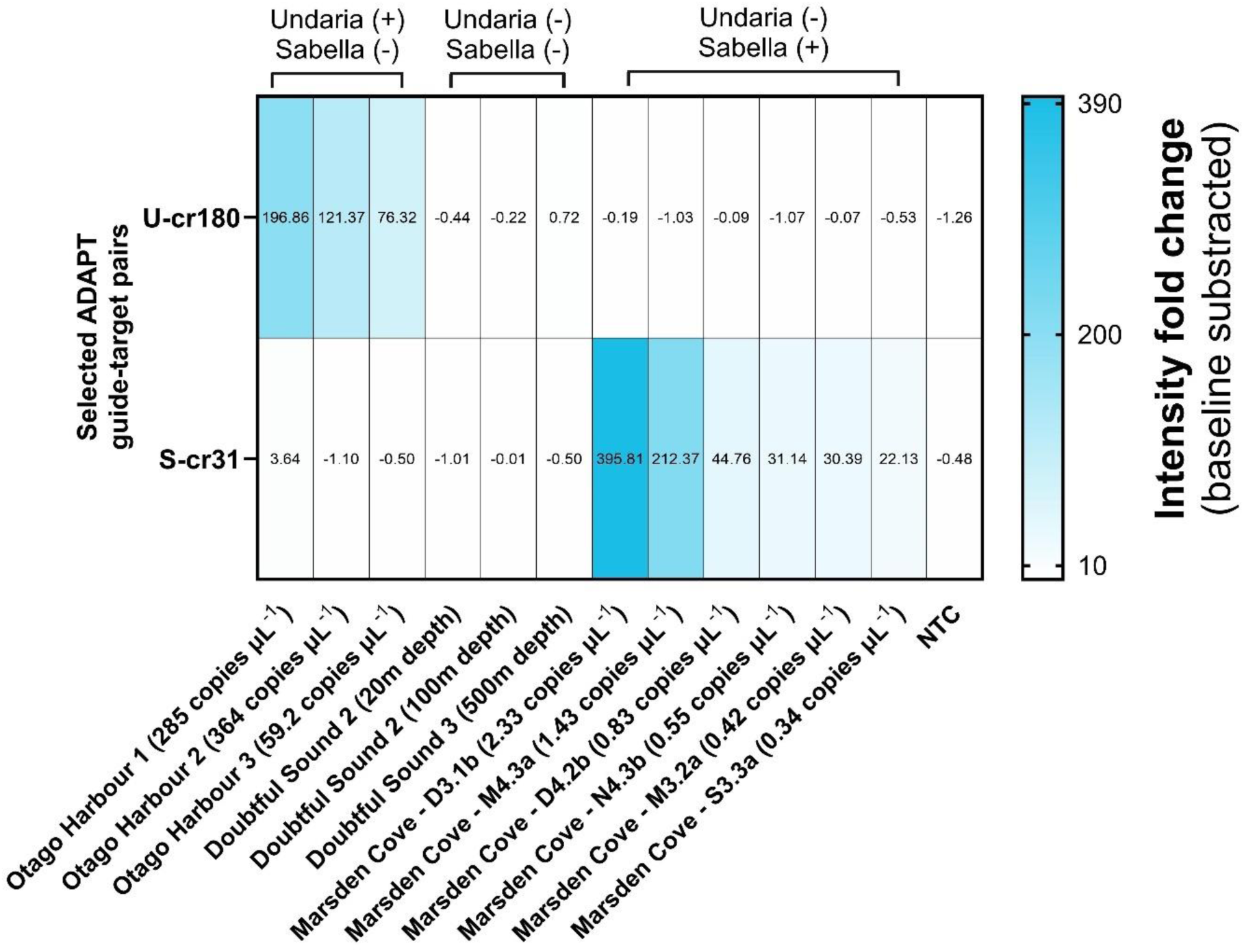
CORSAIR platform in environmental samples. All boxes represent the mean (n = 3). Otago Harbour has a known invasion of *Undaria pinnatifida*, but *Sabella spallanzanii* was not reported when samples were collected. A similar case with *S. spallanzanii* in Marsden Cove, with known *S. spallanzanii* invasion but no reported *U. pinnatifida* invasion. Doubtful sound was used as an environmental DNA negative control free of *S. spallanzanii* and *U. pinnatifida*.

## Discussion

Amid the ongoing global biodiversity crisis in aquatic habitats, driven primarily by climate change and bioinvasions (1), eDNA monitoring holds transformative potential for environmental biosurveillance. Its ability to provide highly accurate species identification, combined with the capacity to survey extensive areas with minimal resource demands, makes it a powerful tool for conservation efforts (4). To address these needs, novel molecular tools are being developed to enhance portability and ease of use without sacrificing specificity and sensitivity in eDNA detection (4, 50, 51). In this context, our study introduces CORSAIR, a new CRISPR-Cas-based strategy designed to detect *S. spallanzanii* and *U. pinnatifida* from eDNA samples.

We first deployed ADAPT locally to design two crRNAs targeting the COI gene regions of *S. spallanzanii* and *U. pinnatifida*, which are known to be species-specific (52, 53). ADAPT identified the shortest possible amplicons (∼88-90 nucleotides per assay) containing highly active guide-target pairs, as illustrated in Figure 2. This feature is particularly advantageous, as RPA performs optimally with smaller amplicons (∼100-200 nucleotides) (54). Additionally, eDNA in natural environments is often fragmented (55), making short amplicons more suitable for detection. By swiftly generating these compact amplicons, ADAPT enhances the efficiency and reliability of eDNA-based biosurveillance.

For the first screening of the CORSAIR platform, we used the guide-target pair U-cr180, where we observed that ADAPT-provided constructs were working as intended, preliminarily highlighting its advantages to streamline LwaCas13a designs for CRISPR-eBx. Moreover, we also observed that the optimized Cas13a reaction buffer yielded the best results, providing an activation speed ∼2.4 times faster than the Cas13a manufacturer buffer (Figure 3). The observed difference in activation speed is likely due to the addition of PEG-8000, which has been reported to enhance enzymatic reaction activity (56, 57). The variation in raw intensity suggests that inactivation of LwaCas13a activity may result from potential depletion of the cofactor (Mg^2+^) rather than a variation of poly-U_5_ reporter, as they were prepared under the same conditions, though the lack of detailed information on the LwaCas13a GenScript reaction buffer limits further analysis. This screening was pivotal for the CORSAIR platform, as slower activation speeds could lead to false negatives, particularly at the single copy level, where reactions are expected to activate more slowly. Therefore, we used the Cas13a optimized buffer for all subsequent CORSAIR assays.

We tested all obtained crRNAs with two different poly-U_x_ reporters (poly-U_5_ and polyU_15_), as longer Uracil chains have been shown to enhance LwaCas13a *trans* activation (47, 58). However, we found that the smaller reporter, poly-U_5_ reported performed better under the same conditions (target concentration: 10 pM final, reporter concentration: 500 nM final), yielding the highest intensity fold change (Figure 4). This discrepancy may be explained by two factors: 1) we employed a novel optimized buffer that accelerates the LwaCas13a reaction (38), whereas other studies used different reaction buffers (27, 43, 53, 54), which can influence reaction kinetics; 2) LwaCas13a *trans*-collateral activity exhibits a dinucleotide preference for uracil pairs (UU) (35), which has been utilized in previous studies using the poly-U_5_ reporter (33, 35, 58).

Assuming that each UU pair results in a single cut, a poly-U_5_ reporter could produce up to two *trans* cuts per reporter molecule. We hypothesize that longer UU chains in a poly-U reporter could lead to more *trans*-cuts, resulting in a similar activation speed (Figure 3) but a lower raw intensity. This may occur because longer poly-U chains could deplete LwaCas13a activity more quickly, potentially due to a finite number of substrate *cis* cuts available per active Cas13a. Hence, *cis*-activated *trans*- collateral activity is limited by the *cis* activity of the Cas13a (59, 60). In perspective, the same molarity was used when comparing poly-U5 and poly-U15 (i.e., the same number of molecules), indicating that in terms of raw intensity release (i.e., required LwaCas13a *trans* cleavage cuts that effectively release the fluorophore), shorter poly-U reporters may be better suited as longer UU chains deplete LwaCas13a activity before the poly-U reporters can provide a maximum intensity. Therefore, from an application perspective, under the conditions of this study, a shorter reporter provides a better activation speed for diagnostics. Consequently, we used poly-U_5_ for all subsequent CORSAIR assays.

Next, we challenged CORSAIR with a proof of concept using off-target species, employing plasmids containing COI inserts from *S. spallanzanii* and *U. pinnatifida*, as well as gDNA from *S. spallazanni*, *U. pinnatifida* and humans. All crRNAs performed as expected, with no detectable off-target activity, even in the presence of potential human gDNA contamination (Figure 5A). Additionally, ADAPT score ranking yielded similar results when comparing crRNAs targeting the same species (e.g., S-cr31 vs S-cr1470) using either plasmid or gDNA (Supplemental Figure 1). Based on these findings, we selected S-cr31 and U-cr180 for all subsequent assays.

In an environmental biosurveillance context, any species of interest will co-exist with other species, which will also be co-captured and co-isolated during eDNA sampling. To account for this, we expanded our off-target assays by creating a panel of 23 potential co-occurring species as potential off-targets for the CORSAIR assay targetting *S. spallanzanii* and *U. pinnatifida* (Figure 5B). No significant intensity fold change was observed above the arbitrary positivity threshold (set at 7-fold-change units, representing the maximum arbitrary background noise).

We then assessed the sensitivity of CORSAIR using plasmids with COI inserts and gDNA from our target species. Our results showed that CORSAIR detected plasmid DNA as low as 10 aM (10^-18^ mol L^-1^ or ∼5-6 copies µL^-1^ or ∼0.024 fg µL^-1^), with a significant endpoint intensity fold change compared to the control (Supplemental Figure 2A and 2B). For gDNA, we detected *S. spallanzanii* at 0.018 pg uL^-1^ and *U. pinnatifida* at 0.036 pg uL^-1^, corresponding to approximately ∼10 fM range (10^-15^ mol L^-1^; Figure 6B), consistent with previous studies using extracted gDNA and LwaCas13a (61). The CORSAIR assay also detected plasmid DNA at 10 aM within 30 minutes (Supplemental Figure 2C), a speed similar to other CRISPR-based assays that outperformed qPCR in terms of result turnaround time (33); however, due to the potential risk of false-negatives, we chose to maintain a 1-hour endpoint to minimize this risk.

Furthermore, both species displayed a semi-quantitative response with both plasmid DNA and gDNA, showing an R^2^ of 0.77 and 0.81 for *S. spallanzanii* and *U. pinnatifida* gDNA samples, respectively. Similar results were obtained with plasmid DNA (Figure 6A).

This semi-quantitative feature of CORSAIR was more pronounced at lower concentrations, as high concentrations quickly saturated the reaction due to faster activation, resulting in the same end-point fluorescence. In contrast, lower-concentration samples did not reach the saturation, leading to varying end-point fluorescence at the 1-hour mark based on concentration, enabling semi-quantification. While this reflects the relative abundance of the eDNA target, it does not directly correlate with species abundance (62) but rather with the genomic region used for detection. Overall, this provided a solid foundation for our CORSAIR assays and gave us confidence to apply them to eDNA samples.

Finally, we tested eDNA samples from locations with known invasions of *S. spallanzanii* (Marsden Cove, New Zealand) and *U. pinnatifida* (Otago Harbour, New Zealand), as well as from a site with known invasions (Doubtful Sound, New Zealand). To validate the CORSAIR assay results, Marsden Cove and Otago Harbour samples were benchmarked against ddPCR and qPCR for *S. spallanzanii* and *U. pinnatifida*, respectively. The result showed 100% agreement with all tested samples: 6/6 for *S. spallanzanii* and 3/3 for *U. pinnatifida*. As expected, no CORSAIR assay exceeded the arbitrary positivity threshold in negative eDNA samples from Doubtful Sound (Figure 7).

When comparing the sensitivity of our CORSAIR assay to that of other recent studies using LwaCas13a (44, 49, 61), similar detection limits were achieved. The best-performing CORSAIR assay detected *S. spallanzanii COI* region at concentrations ranging from 2.33 copies µL^-1^ to 0.34 copies µL^-1^, corresponding to approximately ∼10 aM range, as determined by ddPCR quantification. This is consistent with other CRISPR-eBx studies that have achieved single copy detection (33, 49, 61). For *U. pinnatifida* eDNA samples, the detection range was from 364 copies µL^-1^ to 59 copies µL^- 1^, which corresponds to around the 0.6 fM, according to qPCR quantification.

When comparing CORSAIR-derived fluorescence intensity with ddPCR or qPCR quantification (in copies µL^-1^) from eDNA samples, we observed strong correlations, with R^2^ = 0.84 for *S. spallanzanii* and 0.53 for *U. pinnatifida* (Supplemental Figure 3). These findings highlight two key features for CORSAIR: 1) the semi-quantitative performance is maintained with eDNA samples, and 2) its results align closely with qPCR/ddPCR, underscoring the potential of CRISPR-eBx as a mainstream tool for biosurveillance. Most *S. spallanzanii* and *U. pinnatifida* eDNA samples could also be called positive within 30 minutes, except for sample S3.3a, which required 40-minutes (Supplemental Figure 4). Moreover, the cost per CORSAIR reaction was approximately 3.5 USD (Supplemental Table 9), comparable to reported costs for similar CRISPR-LwaCas13a assays (49).

During this study, a new invasion of *S. spallanzanii* was reported near Otago Harbour (Port Chalmers) between August 26-30, 2024, with a full specimen retrieved (transect coordinates: - 45.81192312, 170.6268429; end of transect:-45.81142622, 170.6271924) However, our assay did not detect this incursion in the Otago Harbour eDNA samples, likely because the invasion occured after our samples were collected on July 17, 2024. Additionally, the sampling site for *U. pinnatifida* was approximately 10.5 km from Port Chalmers. These findings highlight the importance of targeted and timely CRISPR-eBx sampling at sites of interest to ensure the accurate detection of biosecurity threats.

The robustness of CORSAIR was demonstrated through its application to 12 eDNA samples, which were previously benchmarked against ddPCR/qPCR, achieving 100% agreement. This success highlight CORSAIR as a reliable proof-of-concept for the rapid design of CRISPR-based diagnostic technologies using AI tools like ADAPT. Remarkably, the designs in this study were generated within minutes, significantly reducing screening efforts while delivering high-quality results with cost-efficiency at approximately 3.5 USD per sample (Supplemental Table 9). However, this study also identified three key areas requiring further development: 1) further co-occurring species must be tested, but also closely related species for *S. spallanzanii* and *U. pinnatifida* to fully comprehend ADAPT capabilities to discover and detect unique diagnostic regions for environmental biosurveillance deployments. 2) Further eDNA samples must be tested from different sampling sites to assess CORSAIR further. 3) ADAPT was trained to deliver highly active diagnostic of viruses, where eDNA samples can be exceedingly more complex; therefore, there is an inherent potential opportunity to train ADAPT but with high-quality CORSAIR eDNA assays to maximize and tailor ADAPT guide-target pairs specifically for environmental biosurveillance deployments context.

### Concluding remarks

Our study successfully demonstrates the use of a novel blend, using RPA-CRISPR-LwaCas13a and AI modelling, namely CORSAIR, for detecting two invasive species, S. spallanzanii and *U. pinnatifida* using eDNA from aquatic habitats as proof of concept of CRISPR-eBx.

Using ADAPT, a pre-trained end-to-end AI model, we were able to swiftly design and deploy highly active guide-target pairs targeting the COI gene region. We achieved high sensitivity and high specificity with no off-target activity across 23 co-occurring species tested. Our findings indicate that CORSAIR can detect DNA concentrations as low as 10 aM in plasmid DNA, within the fM range in gDNA, and down to 0.32 copies µL^-1^ in eDNA samples. Additionally, we observed a semi-quantitative detection window, allowing relative assessment of eDNA abundance based on endpoint intensity at low concentrations. This feature provides higher confidence in environmental biosurveillance for future applications, where target eDNA is often present in trace amounts. When testing CORSAIR on eDNA samples from two known invasion sites (Marsden Cove and Otago Harbour), it yielded results consistent with ddPCR and qPCR benchmarks for *S. spallanzanii* and *U. pinnatifida*, showcasing a 100% agreement and affirming the robustness and reliability of CORSAIR for environmental biosurveillance with an average cost of ∼3.5 USD per reaction.

However, while CORSAIR shows promise, three key limitations require attention for further research: expanding off-target activity testing on closely related species to ensure specificity, validating across diverse sampling sites, and tailoring ADAPT to optimize guide-target pairs specifically for eDNA contexts, as eDNA samples can have a complex composition of diverse eDNA from difference sources and inhibitors. Another technical development required is to reduce the mixing steps of overall CRISPR-eBx deployments. Addressing these areas will enhance CORSAIR application in real-world environmental biosurveillance efforts worldwide, supporting global biodiversity efforts through rapid and precise CRISPR-eBx.

## Data availability

ADAPT public, open-source repository is available at https://github.com/broadinstitute/adapt

## Supplementary Data

Supplementary data and raw data will be available upon request.

## Acknowledgements

All authors thank the Cawthron Institute (Nelson, New Zealand), National Institute of Water and Atmospheric Research (NIWA), Prof. Joe Zuccarello, Prof. Sarah Caronni and Michelle Liddy for kelp genomic DNA samples. All authors also thank Dr Indranil Basik from the Department of Biochemistry (University of Otago, Dunedin, New Zealand) for human genomic DNA samples.

## Author contributions

**Benjamín Durán Vinet**: conceptualization, methodology, investigation, software, illustration, resources, data curation, formal analysis, writing – original draft and writing – review and editing. **Jo-Ann L. Stanton**: conceptualization (supporting), methodology, investigation, resources, formal analysis, writing – original draft, writing – review and editing and supervision. **Gert-Jan Jeunen:** methodology (supporting), investigation, resources, formal analysis (supporting), writing – review and editing and supervision. **Ulla von Ammon:** investigation (supporting), resources, writing – original draft and writing – review and editing. **Jackson Treece:** methodology (supporting), investigation (supporting) and resources. **Xavier Pochon:** investigation (supporting), resources, writing – review and editing, and supervision. **Anastasija Zaiko:** investigation (supporting), resources, writing – review and editing, supervision, project administration and funding acquisition. **Neil Gemmell:** conceptualization (supporting), methodology (supporting), investigation (supporting), resources, formal analysis, writing – original draft, writing – review and editing, supervision, project administration and funding acquisition.

