## Supplemental for "CRISPR-based environmental biosurveillance assisted via artificial intelligence design of guide-RNAs"

### Supplemental figures

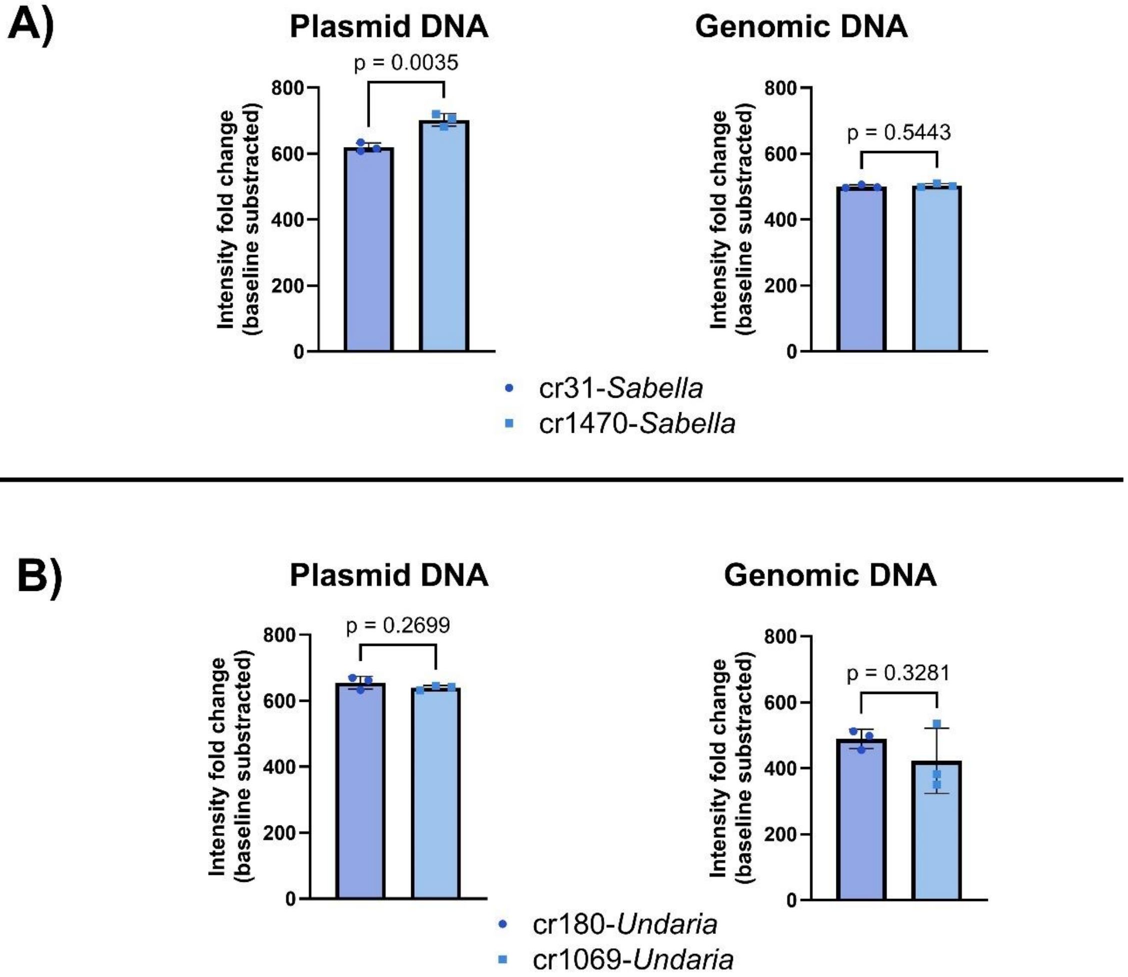


**Supplemental Figure 1. Internal comparison of CORSAIR assay performance using poly-$U_5$ reporter. (A)** Comparison of *S. spallanzanii* guide-target pairs S-cr31 and S-cr1470 when using plasmid and genomic DNA. **(B)** Comparison of *U. pinnatifida* guide-target pairs U-cr180 and U-cr1069 when using plasmid and genomic DNA. Columns represent the mean and error bars show ± SD (n = 3)

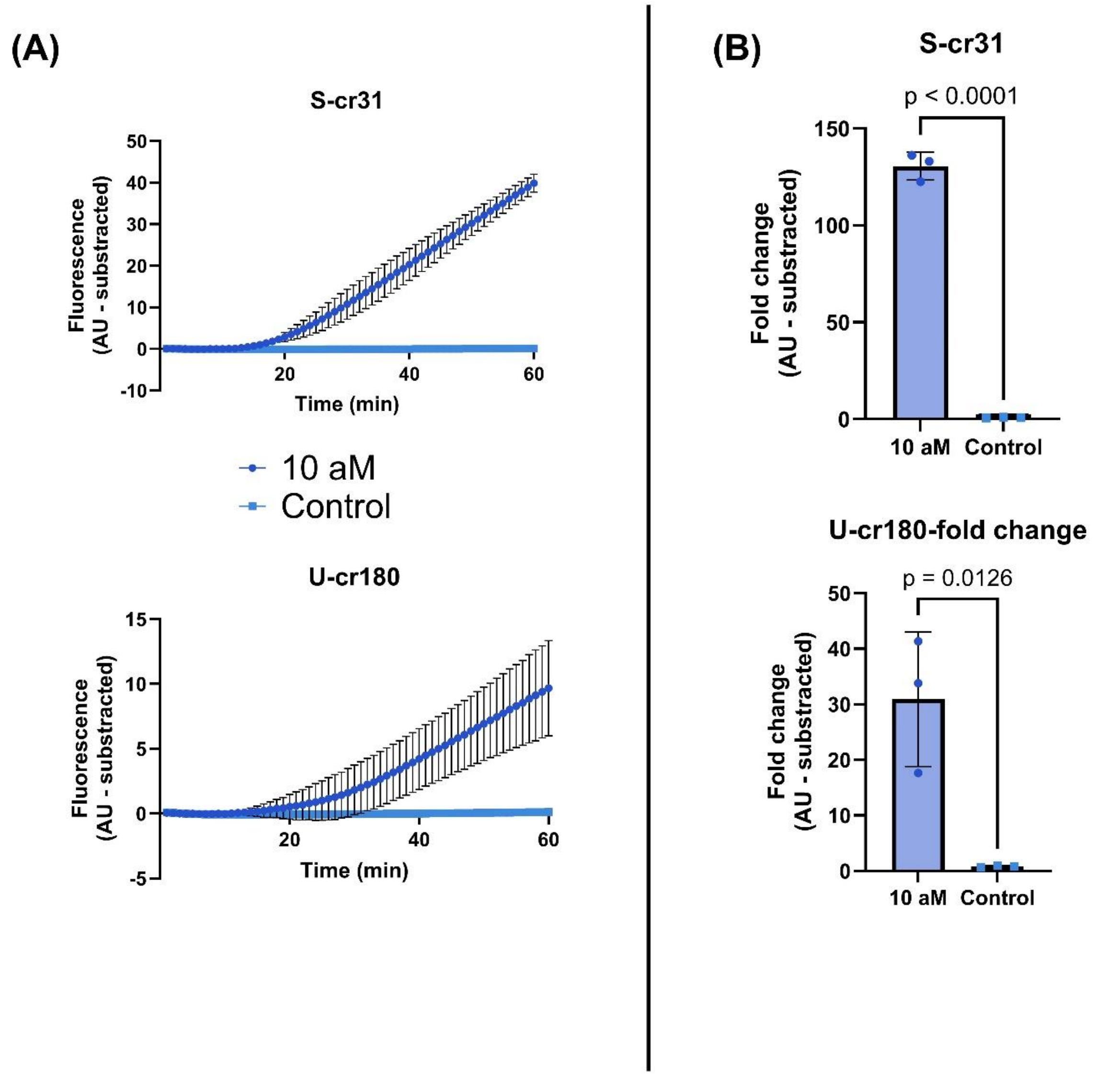


**Supplemental Figure 2. CORSAIR selected guide-target pairs activity at 10 aM range. (A)** Examples of S-cr31 and U-cr180 at 10 aM concentration. **(B)** Significance of intensity fold change at 10 aM versus control ($p<0.05$). Columns represent the mean, and error bars show ± SD (n = 3)

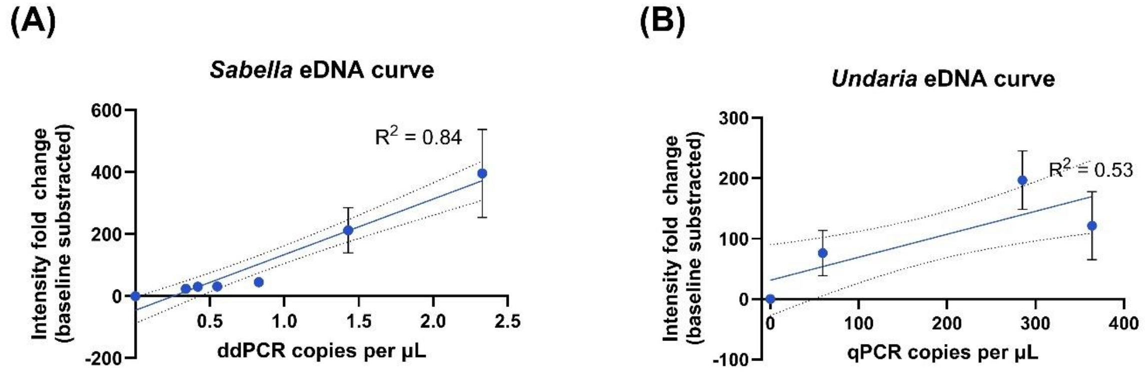


**Supplemental Figure 3. Environmental samples curve fitting when comparing CORSAIR intensity fold change and ddPCR/qPCR quantification. (A)** *Sabella spallanzanii* environmental DNA curve. **(B)** *Undaria pinnatifida* environmental DNA curve. All data points represent the mean. Error bars show ± SD (n = 3). The dotted line represents the 95% confidence interval.

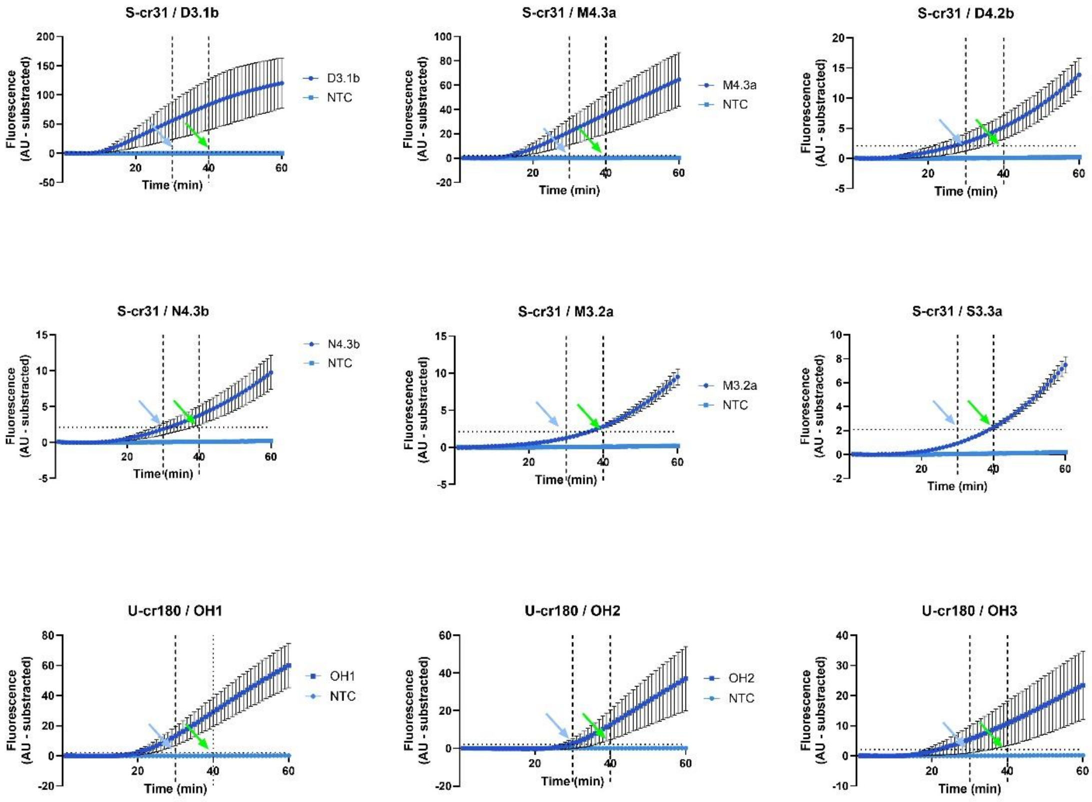


**Supplemental Figure 4. Raw fluorescence intensity plots for all eDNA samples used in this study.** Horizontal dotted line Vertical dotted line shows the arbitrary positive threshold, while vertical lines showcase 30-minute (light blue arrow) and 40-minute mark (green arrow). Error bars show ± SD (n = 3).

### Supplemental tables

**Supplemental Table 1**. Accession numbers for *Sabella spallanzanii* and *Undaria pinnatifida* were used in this study to build consensus sequences.

| Specie name | GenBank accession number (on-target) | GenBank accession number (off-target) |
|---|---|---|
| *Sabella spallanzanii* | KY472735; KY472742; KY472743; KY472759; KY472779; LT717704; LT717705; LT717706; LT717707; LT717708; LT717709; LT717710; LT717711; LT717712; MN184708; MW727509; MZ435261; NC_056279 | LC536448; KR916924; KF369181; AY633070; MG884612 |
| *Undaria pinnatifida* | AB443532; AB443533; AB443534; AB443535; AB443536; AB775223; AF037993; EF218853; GQ368267; KC782871; KY047176; KY047177; KY682973; NC_023354 | HQ848896; KT315643; LC148150; LC515364; NC_013476; OK149005; OK149033; OK149037; OL456970 |

**Supplemental Table 2. Consensus sequence for plasmid insert.**

| Name | Sequence (5’-3’) |
|---|---|
| *Undaria pinnatifida* COI insert | GGTACATTATATTTAATTTTTGGGGGTTTTTCGGGGGTTCTTGGTACAGCAATGT<br>CTGTTTTTATCCGATTGCAATTAGCTAGCCCTGGTAATCAATTTTTGGGGGGAAA<br>TCATCAGTTGTATAATGTTATTGTAACAGCACATGCTTTCTTAATGATTTTTTTTATG<br>GTTATGCCAATTCTTATCGGTGGATTTGGTAACTGGTTTGTACCTTTAATGATTGG<br>TGCTCCTGATATGGCTTTTCCTCGTATGAATAATATTAGTTTTTGGTTATTACCTCC<br>CTCTTTAATTCTTCTTTTAGCCTCTTCTTTAGTAGAGTCTGGGGCTGGAACAGGT<br>TGGACGGTATACCCTCCGCTTAGTGGTATTCAAGCTCACTCAGGTCCTTCAGT<br>TGATTTAGCTATTTTTAGTCTTCACCTTTCAGGAGCTGCTTCTATTTTAGGTGCTAT<br>AAACTTTATTACCACAATTTTTAATATGAGAGCACCTGGTATGACAATGGATAGAT<br>TGCCTCTTTTTGTGTGGTCTGTTTTAATTACAGCGTTTTTATTGTTGTTATCATTGC<br>CGGTTTTAGCAGGTGCTGTTACAATGTTACTAACAGATCGTAATTTTAATACTACT<br>TTTTTTGATCCTGCGGGCGGTGGTGATCCAGTATTATATCAGCATTTATTTTGGTT<br>CTTTGGTCATCCTGAAGTATATATATTAATTTTACCAGGATTCGGTATTGTTAGTCA<br>TATATTATCTACTTTATCAAGAAAACCAGTTTTCGGTTATTTAGGTATGGTTTATGC<br>TATGCTTTCTATAGGTATACTTGGTTTTATTGTATGGGCTCATCACATGTTTACAGT<br>AGGTTTAGATATTGATACAAGAGCTTATTTTACAGCAGCCACTATGATTATTGCG<br>GTCCCTACAGGTATTAAAATTTTTAGTTGGGTCGCAACTTTGTGGGGAGGTTCT<br>ATTCGGTTAAAAACTCCGATGTTATTTTCTATAGGTTTTCTTTTTCTTTTTACTATAG<br>GAGGGTTAACAGGGGGTTGTTTTAGCAAATTCAGGGGGTTGATATTGCTTTA |
| *Sabella spallanzanii* COI insert | ATTTTCTCTCGGTGGATATGGTCTAGGAATCACAAGGATATTGGAACTTTGTATTT<br>TATTTTGGGTTTGTGAGGTGGGCTTGTTGGAACTTCTATAAGAATTTTAATTCGTTT<br>GGAGTTAGGTCAGCCTGGAACTTTGTTGGGAAGTGATCAGTTGTATAATTCTATT<br>GTAACTGCCCATGCATTTTTAATAATTTTTTTTTTAGTTATGCCTGTTTTTATTGGTG<br>GATTTGGAAATTGATTATTGCCCCTGATGATTGGGGCCCCAGATATAAGATTCC<br>CCCGGCTTAATAATCTAAGGTTTTGACTACTACCTTCTGCATTATTGCTCTTATTA<br>GGCTCTGTGTTTGTAGAAACTGGGGCAGGGACGGGATGAACTATTTATCCTCC<br>TTTGTCAAGAGGAGTTGGACATAGAGGGTCTTCCATGGATTTAGTAATTTTTTCA<br>CTTCATTTGGCAGGGGCTTCTTCTATTATAGGGGCAGTAAACTTTATTACTACTAT<br>TGCTAATTTAGGATCTAGTTCTATACGGGGAGAGCGCTTACCTTTATTTGTTTGA<br>GCAGTAGTTATTACTGTTGTTTTATTATTATTGTCTTTGCCTGTGTTGGCGGCGGC<br>TATTACTATATTATTAACAGATCGAAATATTAATACAAGGTTTTTTGACCCATCTGG<br>TGGAGGAGATCCGGTATTATTTCAGCATTTATTTTGGTTTTTTGGTCATCCTGAG |

| | |
|---|---|
| | GTTTATATTTTAATTCTTCCTGCATTTGGGGTGGTATCCCATGTTGTTACCCATTTT<br>TCTGGGAAGTTAAGTTCTTTTGGTTTGCTTGGTATAATTTATGCTATACTAAGTATT<br>GGGGTATTAGGGTTTCTTGTTTGGGCACACCATATATTTACAGTTGGAATGGATG<br>TTGATACTCGGGCATATTTTACTTCTGCTACTATAACTATTGCTGTTCCAACGGG<br>GATTAAGGTGTTTAGCTGACTTGCTACCTTAAGTGGGGGAAAAGTAAGGTGTGA<br>GGGTCCTTTGTTATGGGTGCTGGGTTTTATTTTTCTCTTTACTGTAGGGGGCTTA<br>ACTGGAGTTATTTTAGCAAATTCTTCTTTAGATATTGCATTACATGACACATATTAT<br>GTGGTGGCTCATTTCCATTATGTGTTGAGAATGGGAGCTATTTTTGGGATTTTTG<br>CAGGGATTACATTTTGATTTCCATTATTTACTGGTGTAGGGTTGCATGGGCGGT<br>GAGTTAAGGGGCATTTTTTCTTAATATTTATTGGTGTTAATTTAACTTTTTTTCCTCA<br>GCATTTCTTAGGGTTAGCCGGAATGCCCCGCCGATATTCTGATTATCCTGATGT<br>ATTTATATCTTGGAATGTTGTATCATCAATTGGGTCTATAATTTCTTTGGTTGGTTTA<br>TTGTTTTTTATATTTATTTTATGGGAGGCTTTGGCGGCTCAGCGTAAATTGCTGTA<br>CACTCCTAGGGCAGGGACAAGATTAGAGTGAAGATGGGCTAGATACCCACTA<br>TCAGCACATACGCACATAGAAGTGGTTATAACTTTTAGT |

Consensus sequences were obtained from aligned sequences from the corresponding species cytochrome c oxidase subunit I (COI) mentioned in Supplemental Table 1.

**Supplemental Table 3. Final code to run ADAPT for *Sabella spallanzanii* and *Undaria pinnatifida*.**

| Specie | Code |
|---|---|
| *Sabella spallanzanii* | design.py complete-targets fasta /Users/durbe957/Desktop/ADAPT_data/Input/Sabella_COI.fasta -o /Users/durbe957/Desktop/ADAPT_data/Output/Sabella_COIfilter-2 --specific-against-fastas /Users/durbe957/Desktop/ADAPT_data/Input/Sabella_Off1.fasta --id-m 2 --id-frac 0.05 --obj maximize-activity -gl 28 -gm 0 -pt -pl 30 -pm 1 -pp 0.95 --penalty-strength 0.25 --primer-gc-content-bounds 0.30 0.55 --maximization-algorithm random-greedy --predict-cas13a-activity-model --best-n-targets 20 --seed 027 --verbose |
| *Undaria pinnatifida* | design.py complete-targets fasta /Users/durbe957/Desktop/ADAPT_data/Input/Undaria_COI.fasta -o /Users/durbe957/Desktop/ADAPT_data/Output/Undaria_COIfilter-2 --specific-against-fastas /Users/durbe957/Desktop/ADAPT_data/Input/Undaria_Off1.fasta --id-m 2 --id-frac 0.05 --obj maximize-activity -gl 28 -gm 0 -pt -pl 30 -pm 1 -pp 0.95 --penalty-strength 0.25 --primer-gc-content-bounds 0.30 0.55 --maximization-algorithm random-greedy --predict-cas13a-activity-model --best-n-targets 20 --seed 026 --verbose |

Further information on the code can be found at GitHub (github.com/broadinstitute/adapt)and also on Metsky et al. (1)

**Supplemental Table 4. Species-specific primers, CRISPR RNAs and probes used in this study.**

| Name | Sequences (5'-3')[a] | Objective score | Amplicon size |
|---|---|---|---|
| **S-31-t7F** | **gaaatTAATACGACTCACTATAGGG**TTTTCTCTCGGTGGATATGGTCTAGGAATC | 6.04 (worst) | 88 bp |
| **S-cr31** | GAUUUAGACUACCCCAAAAACGAAGGGGACUAAAACAUAAAAUACAAAGUUCCAAUAUCCUUGU | | |
| **S-31-R** | GTTCCAACAAGCCCACCTCACAAACCCAAA | | |
| **S-1470-t7F** | **gaaatTAATACGACTCACTATAGGG**TTGCTGTACACTCCTAGGGCAGGGACAAGA | 6.34 (best) | 88 bp |

| | | | |
|---|---|---|---|
| **S-cr1470** | GAUUUAGACUACCCCAAAAACGAAGGGGACUAAAACGUGGGUAUCUAGCCCAUCUUCACUCUAA | | |
| **S-1470-R** | TAACCACTTCTATGTGCGTATGTGCTGATA | | |
| **U-180-t7F** | **gaaatTAATACGACTCACTATAGGG**GGGGTTCTTGGTACAGCAATGTCTGTTTTT | | |
| **U-cr180** | GAUUUAGACUACCCCAAAAACGAAGGGGACUAAAACUACCAGGGCUAGCUAAUUGCAAUCGGAU | 6.11 (worst) | 88bp |
| **U-180-R** | ACAACTGATGATTTCCCCCCAAAAATTGAT | | |
| **U-1069-t7F** | **gaaatTAATACGACTCACTATAGGG**CAGGTATTAAAATTTTTAGTTGGGTCGCAA | | |
| **U-cr1069** | GAUUUAGACUACCCCAAAAACGAAGGGGACUAAAACGAAAAUAACAUCGGAGUUUUUAACCGAA | 6.13 (best) | 110bp |
| **U-1069-R** | CCCTCCTATAGTAAAAAGAAAAAGAAAACC | | |
| **U5-reporter** | 6-FAM/UUUUU/BHQ-1 | N/A | N/A |
| **U15-reporter** | 6-FAM/UUUUUUUUUUUUUUU/BHQ-1 | N/A | N/A |

[a] Direct repeat region is underlined in CRISPR RNA (crRNAs) sequences. 5' T7 promoter is bolded in forward primers. The spacer region is left unmodified in the crRNAs.

### Supplemental Table 5. CORSAIR mastermix setup.

**Table 2.** Cas13 mastermix detailed volumetric composition.

| *Component* | **20 μL reaction** | **4 reactions[a]** |
|---|---|---|
| RNAse-free water | 5.18 | 43.51 |
| Reaction buffer (5X)[c] | 4.00 | 33.60 |
| Murine RNAse inhibitor (40 U/μL) | 0.50 | 4.20 |
| rNTP mix (25 mM) | 0.80 | 6.72 |
| Reporter (10 μM) | 1.00 | 8.40 |
| Forward primer (10 μM) | 0.96 | 8.06 |
| Reverse primer (10 μM) | 0.96 | 8.06 |
| T7 RNA polymerase (50 U/μL) | 0.6 | 5.04 |
| LwaCas13a (500 nM) | 2.00 | 8.40 |
| crRNA (1 μM) | 1.00 | 4.20 |
| MgOAc (280 mM) | 1.00 | 4.20 |
| **RPA pellet** | **N/A** | **1** |
| **Total to resuspend** | **N/A** | **75.60** |
| **Aliquot size** | **N/A** | **18** |
| **Sample[b]** | **N/A** | **2** |
| **Total wells** | **N/A** | **4** |

N/A: Not applicable.

[a] 5% pipetting error was introduced into the calculation.

[b] The sample is added after aliquoting the 18 uL of the mastermix in each well of the strip/plate.

[c] Reaction buffer composition is available in Supplemental Table 6.

### Supplemental Table 6. Reaction buffer composition.

| **Component** | **Supplier** | **Final 5X concentration** |
|---|---|---|
| **HEPES 1M pH 8.0** | ThermoFisher | 100 mM |
| **KCl 2M** | ThermoFisher | 300 mM |
| **PEG-8000** | Merck (Sigma Aldrich) | 25% m/v |

Ensure all components are molecular grade.

Buffer components were obtained from Arizti-Sanz et al. (2).

**Supplemental Table 7. ADAPT guide-target pair, position, and objective scores obtained for *Sabella spallanzanii* and *Undaria pinnatifida*.**

| Specie | Position | Objective score[a] | Forward primer | Reverse[b] | Spacer[b] |
|---|---|---|---|---|---|
| **Sabella spallanzanii** | {1470} | **6.34** | TTG CTG TAC ACT CCT AGG GCA GGG ACA AGA | TAT CAG CAC ATA CGC ACA TAG AAG TGG TTA | TTA GAG TGA AGA TGG GCT AGA TAC CCA C |
| | {1156} | 6.27 | TCC ATT ATG TGT TGA GAA TGG GAG CTA TTT | ATT TCC ATT ATT TAC TGG TGT AGG GTT GCA | TTG GGA TTT TTG CAG GGA TTA CAT TTT G |
| | {1327} | 6.22 | CCG GAA TGC CCC GCC GAT ATT CTG ATT ATC | ATC ATC AAT TGG GTC TAT AAT TTC TTT GGT | CTG ATG TAT TTA TAT CTT GGA ATG TTG T |
| | {836} | 6.18 | GGT TTG CTT GGT ATA ATT TAT GCT ATA CTA | CAT ATA TTT ACA GTT GGA ATG GAT GTT GAT | TGG GGT ATT AGG GTT TCT TGT TTG GGC A |
| | {943} | 6.14 | CTT CTG CTA CTA TAA CTA TTG CTG TTC CAA | TAC CTT AAG TGG GGG AAA AGT AAG GTG TGA | CGG GGA TTA AGG TGT TTA GCT GAC TTG C |
| | {31} | **6.04** | TTT TCT CTC GGT GGA TAT GGT CTA GGA ATC | TTT GGG TTT GTG AGG TGG GCT TGT TGG AAC | ACA AGG ATA TTG GAA CTT TGT ATT TTA T |
| **Undaria pinnatifida** | {1069} | **6.13** | CAG GTA TTA AAA TTT TTA GTT GGG TCG CAA | GGT TTT CTT TTT CTT TTT ACT ATA GGA GGG | TTC GGT TAA AAA CTC CGA TGT TAT TTT C |
| | {1472} | 6.19 | TGG TTC AAT TTT ATC TTC TAT CGC ATC ATT | CTG ACA AGA GGT TCT ATT GAA GAA TCA AAT | ATT TTT TTT TTT TGT TGT TTA TAT TAC T |
| | {849} | 6.18 | TTA CCA GGA TTC GGT ATT GTT AGT CAT ATA | TCG GTT ATT TAG GTA TGG TTT ATG CTA TGC | TTA TCT ACT TTA TCA AGA AAA CCA GTT T |
| | {180} | **6.11** | GGG GTT CTT GGT ACA GCA ATG TCT GTT TTT | ATC AAT TTT TGG GGG GAA ATC ATC AGT TGT | ATC CGA TTG CAA TTA GCT AGC CCT GGT A |

[a] Best and worst scores for each species are bolded.

[b] Reverse and spacer sequences must be the reverse complement to be functional.

**Supplemental Table 8**. Samples used in this study.

| Source | Assay | ID | Coordinates/Source | Status | Reference |
|---|---|---|---|---|---|
| **Marsden Cove** | CORSAIR eDNA | D3.1b | -35.836802, 120 174.468679 *Collected between the 19-06-21 and 20-06-21* | Sabella positive (qPCR) | (3) |
| | CORSAIR eDNA | M4.3a | -35.836802, 120 174.468679 | Sabella positive (qPCR) | (3) |

| | | | | | |
|---|---|---|---|---|---|
| | | | *Collected between the 19-06-21 and 20-06-21* | | |
| | CORSAIR eDNA | D4.2b | -35.836802, 120 174.468679<br>*Collected between the 19-06-21 and 20-06-21* | Sabella positive (qPCR) | (3) |
| | CORSAIR eDNA | N4.3b | -35.836802, 120 174.468679<br>*Collected between the 19-06-21 and 20-06-21* | Undaria negative (qPCR) | (3) |
| | CORSAIR eDNA | M3.2a | -35.836802, 120 174.468679<br>*Collected between the 19-06-21 and 20-06-21* | Undaria negative (qPCR) | (3) |
| | CORSAIR eDNA | S3.3a | -35.836802, 120 174.468679<br>*Collected between the 19-06-21 and 20-06-21* | *Undaria* negative (qPCR) | (3) |
| **Tissue extractions** | gDNA testing | S-3 | N/A | N/A | This study |
| | gDNA testing | U-3 | N/A | N/A | This study |
| **Otago Harbour, New Zealand** | CORSAIR eDNA | OH1 | -45.827979, 170.639868 –<br>*Collected on the 17-07-24* | *Undaria* sighted, qPCR positive (see Supplemental Data 2) | This study |
| | CORSAIR eDNA | OH2 | -45.827979, 170.639868<br>*Collected on the 17-07-24* | *Undaria* sighted, qPCR positive (see Supplemental Data 2) | This study |
| | CORSAIR eDNA | OH3 | -45.827979, 170.639868<br>*Collected on the 17-07-24* | *Undaria* sighted, qPCR positive (see Supplemental Data 2) | This study |
| **Doubtful sound, New Zealand** | CORSAIR eDNA | D0 | See reference | *Undaria/Sabella* negative | (4) |
| | CORSAIR eDNA | D4 | See reference | *Undaria/Sabella* negative | (4) |
| | CORSAIR eDNA | D15 | See reference | *Undaria/Sabella* negative | (4) |
| *Cystophora torulosa* | CORSAIR off-target assay | CKK-1 | Kaka Point, Catlins (Cawthron Institute, New Zealand) | N/A | This study |

| | | | | | |
|---|---|---|---|---|---|
| *Chondria macrocarpa* | CORSAIR off-target assay | CKK-2 | Kaka Point, Catlins (Cawthron Institute, New Zealand) | N/A | This study |
| Unidentified red algae | CORSAIR off-target assay | CKK-3 | Kaka Point, Catlins (Cawthron Institute, New Zealand) | N/A | This study |
| *Scytothamnus australis* | CORSAIR off-target assay | CKK-4 | Kaka Point, Catlins (Cawthron Institute, New Zealand) | N/A | This study |
| *Xiphophora gladiata* | CORSAIR off-target assay | CKK-5 | Kaka Point, Catlins (Cawthron Institute, New Zealand) | N/A | This study |
| *Codium fragile* | CORSAIR off-target assay | CKK-6 | Kaka Point, Catlins (Cawthron Institute, New Zealand) | N/A | This study |
| *Cystophora retroflexa* | CORSAIR off-target assay | CKK-7 | Kaka Point, Catlins (Cawthron Institute, New Zealand) | N/A | This study |
| *Macrocystis pyrifera* | CORSAIR off-target assay | CKK-8 | Kaka Point, Catlins (Cawthron Institute, New Zealand) | N/A | This study |
| *Durvillaea poha* | CORSAIR off-target assay | CKK-9 | Kaka Point, Catlins (Cawthron Institute, New Zealand) | N/A | This study |
| *Bryopsis vestita* | CORSAIR off-target assay | Bry10-1 | St. Kilda, Otago (New Zealand) | N/A | This study |
| *Caulerpa taxifolia* | CORSAIR off-target assay | C.tax-1 | Sardinia (Italy) | N/A | This study |
| *Caulerpa proliferia* | CORSAIR off-target assay | C.prolif-1 | Sardinia (Italy) | N/A | This study |
| *Caulerpa brownii* | CORSAIR off-target assay | C.brown-1 | St. Kilda, Otago (New Zealand) | N/A | This study |
| *Caulerpa brachypus* | CORSAIR off-target assay | C1-1 | Great Barrier Island (New Zealand) | N/A | This study |
| *Caulerpa parvifolia* | CORSAIR off-target assay | C3-1 | Great Barrier Island (New Zealand) | N/A | This study |
| *Caulerpa cylindracea* | CORSAIR off-target assay | A25 | Wellington Pet Shop (New Zealand) | N/A | This study |
| *Caulerpa lentillifera* | CORSAIR off-target assay | A26 | Wellington Pet Shop (New Zealand) | N/A | This study |
| *Caulerpa selago* | CORSAIR off-target assay | A40 | Wellington Pet Shop (New Zealand) | N/A | This study |

| | | | | | |
|---|---|---|---|---|---|
| *Caulerpa serrulata* | CORSAIR off-target assay | A45 | Wellington Pet Shop (New Zealand) | N/A | This study |
| *Caulerpa sertularioides* | CORSAIR off-target assay | A78 | Wellington Pet Shop (New Zealand) | N/A | This study |
| *Caulerpa racemosa* | CORSAIR off-target assay | A79 | Wellington Pet Shop (New Zealand) | N/A | This study |
| *Caulerpa chemnitzia* | CORSAIR off-target assay | A91 | Wellington Pet Shop (New Zealand) | N/A | This study |
| *Caulerpa nummularia* | CORSAIR off-target assay | A111 | Wellington Pet Shop (New Zealand) | N/A | This study |

**Supplemental Table 9. CORSAIR cost breakdown.**

| **Component** | ***Supplier*** | ***Quantity*** | ***Net reactions*** | ***Unit value (USD)*** | ***Value per reaction (USD)*** |
|---|---|---|---|---|---|
| **LwaCas13a** | GenScript | 100 ug | 713 | 189 | **0.27** |
| **Forward primer** | GenScript | 20 nmol | 833 | 11.55 | **0.01** |
| **Reverse primer** | GenScript | 20 nmol | 833 | 6.3 | **0.01** |
| **Guide RNA** | GenScript | 4 nmol | 833 | 224 | **0.27** |
| **Cas13-reporter** | GenScript | 12 nmol | 1200 | 165 | **0.14** |
| **RPA kit** | TwistDx | 96 reactions | 384* | 443 | **1.15** |
| **rNTP mix** | NEB | 50 umol | 2000 | 472.08 | **0.24** |
| **Murine RNAse inhibitor** | NEB | 40,000 U/mL | 150 | 114.83 | **0.77** |
| **T7 polymerase** | NEB | 50,000 U/mL | 167 | 105 | **0.63** |
| | | | | **Cost per reaction** | **3.5** |

*Each RPA pellet yields ~4 reactions.
